# Dynamic rewiring of the human interactome by interferon signalling

**DOI:** 10.1101/766808

**Authors:** Craig H. Kerr, Michael A. Skinnider, Angel M. Madero, Daniel D.T. Andrews, R. Greg Stacey, Queenie W.T. Chan, Nikolay Stoynov, Eric Jan, Leonard J. Foster

## Abstract

**Background:** The type I interferon (IFN) response is an ancient pathway that protects cells against viral pathogens by inducing the transcription of hundreds of IFN-stimulated genes (ISGs). Transcriptomic and biochemical approaches have established comprehensive catalogues of ISGs across species and cell types, but their antiviral mechanisms remain incompletely characterized. Here, we apply a combination of quantitative proteomic approaches to delineate the effects of IFN signalling on the human proteome, culminating in the use of protein correlation profiling to map IFN-induced rearrangements in the human protein-protein interaction network.

**Results:** We identified >27,000 protein interactions in IFN-stimulated and unstimulated cells, many of which involve proteins associated with human disease and are observed exclusively within the IFN-stimulated network. Differential network analysis reveals interaction rewiring across a surprisingly broad spectrum of cellular pathways in the antiviral response. We identify IFN-dependent protein-protein interactions mediating novel regulatory mechanisms at the transcriptional and translational levels, with one such interaction modulating the transcriptional activity of STAT1. Moreover, we reveal IFN-dependent changes in ribosomal composition that act to buffer ISG protein synthesis.

**Conclusions:** Our map of the IFN interactome provides a global view of the complex cellular networks activated during the antiviral response, placing ISGs in a functional context, and serves as a framework to understand how these networks are dysregulated in autoimmune or inflammatory disease.

## BACKGROUND

Type I interferons (IFNs) are an evolutionary ancient family of cytokines that play a central role in the immune response to viral pathogens [1]. IFN synthesis and secretion is triggered in response to pathogen detection by intra- and extracellular receptors, leading to the activation of multiple defense mechanisms via the transcription of IFN-stimulated genes (ISGs) [2]. These ISGs contribute to the establishment of a cell-intrinsic antiviral state in infected and neighboring cells, while also modulating the development of innate and adaptive immune responses [3]. Activation of the IFN response must be carefully regulated in order to strike a balance between effective pathogen clearance on the one hand and tissue damage or auto-inflammatory pathology on the other, as aberrant IFN signalling has been implicated in a range of autoimmune and neuropsychiatric diseases [4, 5].

In the canonical type I IFN signalling pathway, IFNs bind the heterodimeric IFNɑ receptor (IFNAR) complex, thereby activating the receptor-associated tyrosine kinases JAK1 and TYK2. In turn, these kinases phosphorylate the cytoplasmic STAT1 and STAT2 transcription factors. Translocation of STAT1 and STAT2 to the nucleus, followed by association with IRF9 to form the IFN-stimulated gene factor 3 (ISGF3) complex, activates ISG transcription. While some of these ISGs encode proteins with direct antiviral activity, many ISG products modulate parallel signalling pathways, or encode additional transcription factors. Consequently, IFN stimulation induces a complex response that is not limited to a simple antiviral program, but instead activates a number of additional signalling pathways such as the MAPK cascade and the mTOR-AKT-S6K axis, which contribute to ISG induction or the antiviral response more broadly [3]. Ultimately, this cascade results in substantial remodelling of mRNA processing, post-translational modification, metabolism, cellular trafficking, chromatin organization, and the cytoskeleton, among other processes [6].

A combination of unbiased transcriptome profiling [2, 7–9] and biochemical approaches [10–13], have identified hundreds of ISGs and, in some cases, elucidated their mechanism of action. Yet the functional roles of most ISGs as effectors of the innate immune response remain to be fully characterized. Furthermore, in view of the limited ability of mRNA levels to predict cellular protein abundance [14, 15], the degree to which IFN-induced changes in transcriptional activity ultimately manifest at the level of the proteome remains incompletely understood. A complete understanding of the IFN signalling repertoire would include a direct interrogation of the complex network of interacting proteins that mediate the type I IFN response, beyond those with a direct role in restricting viral replication. However, experimentally mapping the cellular interaction network in differential and physiologically relevant contexts at the proteome scale represents a longstanding challenge [16].

Here, we apply a combination of quantitative proteomic approaches to chart the molecular landscape of type I IFN signalling, culminating in the use of protein correlation profiling (PCP) [17] to map interferon-induced rearrangements in the human interactome. The resulting protein-protein interaction network, encompassing over 27,000 interactions, reveals widespread rewiring of physical interactions and places known ISGs in an IFN-dependent functional context. We find evidence that the most evolutionarily conserved subset of ISGs are induced to physically interact in response to IFN stimulation, and experimentally validate the role of one such interaction in modulating STAT1 transcription. We develop statistical methods for differential network analysis to characterize interactome rewiring at the functional level, leading us to identify alterations in ribosome composition induced by interferon signalling that selectively downregulate ISG synthesis in order to fine-tune the IFN response. Collectively, this differential network map of the IFN-induced interactome provides a resource to mechanistically dissect the IFN response in the context of viral infection and autoimmune disease.

## RESULTS

### Proteome-wide analysis of the type I IFN response

Whereas the transcriptional response to IFN stimulation has been extensively characterized, the dynamic changes occuring at the proteome level remain unclear. In view of the multiple biological mechanisms that exist to decouple protein abundance from mRNA expression [18], we therefore first sought to establish the proteome-wide response to IFN stimulation. We applied stable isotopic labeling by amino acids in cell culture (SILAC)-based mass spectrometry to precisely quantify protein abundance in HeLa cells after 4 h or 24 h of IFN stimulation (Figure 1A). A total of 7,421 proteins were identified, of which 5,016 were quantified in all three replicates (Table S1). After 4 h of IFN stimulation, a time point by which most ISGs have reached their maximal expression [8], we detected 924 differentially expressed proteins at a 5% FDR, but only 36 with greater than two-fold induction (Figure 1B). Conversely, after 24 h of IFN stimulation, we observed more pronounced changes in the cellular proteome, with 1,172 proteins differentially expressed at 5% FDR and 105 with at least a two-fold induction (Figure 1C and Figure S1A). Several proteins with well-appreciated roles in the type I IFN response were induced over 100-fold, including the IFIT proteins (IFIT1, IFIT2, and IFIT3), MX1, and ISG15 (Table S1).

**Figure 1.**
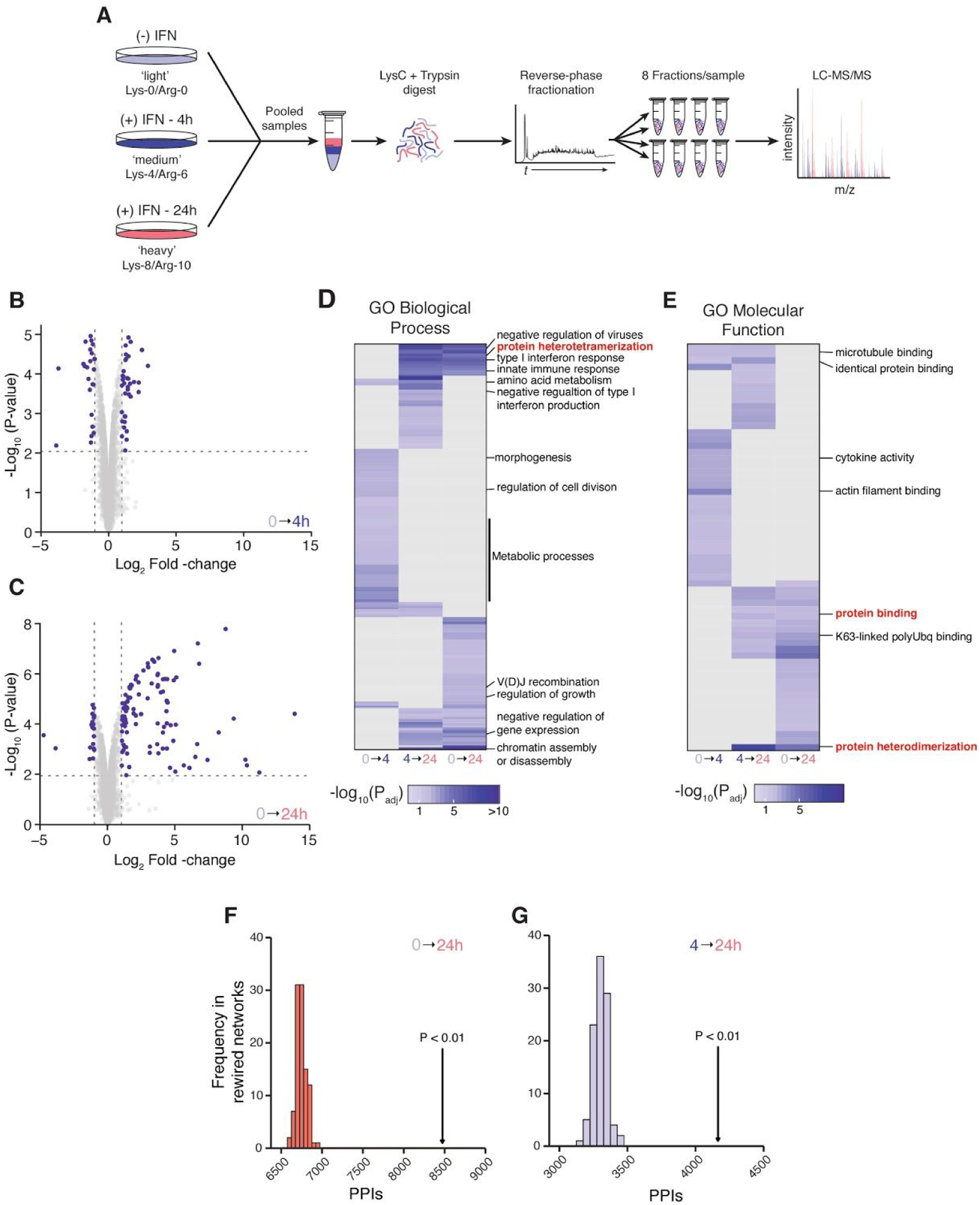
Quantitative proteomic analysis of IFNβ stimulation. (A) Schematic overview of the shotgun proteomics workflow for analysis of IFN-induced proteome changes. (B–C) Volcano plot showing differential protein abundance in cells stimulated with IFNβ for 4 h (B) or 24 h (C). Vertical lines denote absolute fold change ≥ 2. Horizontal lines show 5% FDR threshold. (D–E) Gene Ontology (GO) terms for biological processes (D) and molecular functions (E) significantly enriched among differentially expressed proteins after 4 h or 24 h of IFNβ stimulation. (F–G) Number of protein-protein interactions in the InBioMap database [22] between differentially expressed proteins after 24 h (F) of IFNβ stimulation, or between cells stimulated for 4 and 24 h (G), arrows, and in 100 randomly rewired networks derived from the same database, histograms.

Functional enrichment analysis of the proteins that were differentially expressed at 4 h or 24 h, or which were differentially expressed between the two timepoints, revealed marked temporal differences in the cellular processes activated by IFN stimulation (Figure 1D–E, Figure S1B and Table S2). Proteins that were differentially expressed at 4 h were enriched for Gene Ontology (GO) terms related to involvement in metabolic processes, consistent with the notion that IFN signalling induces changes to cellular metabolism in order to establish an antiviral state [19, 20]. In contrast, the most significantly enriched GO terms at 24 h were related to chromatin rearrangements, in line with the finding that IFN stimulation induces chromatin modifications to establish a transcriptional ‘memory’, resulting in faster and greater transcriptional responses upon restimulation [21]. Surprisingly, despite the rapid induction of ISG transcription (as early as 30 min post-stimulation; [8]), we did not observe an enrichment of Gene Ontology (GO) terms related to the innate immune response until 24 h, reflecting the apparent lag in translation of canonical ISGs (Figure 1B–C).

Our attention was drawn to the enrichment for GO terms related to protein-protein interactions after 24 h of IFN stimulation, including “protein binding,” “protein heterodimerization,” and “protein heterotetramerization.” We therefore sought to determine whether proteins that were differentially expressed at 24 h, relative to 4 h or to unstimulated cells, displayed a statistically significant tendency to physically interact. Comparing the observed number of protein-protein interactions in the InBio Map database [22] to randomly rewired networks [23] revealed a substantial excess of physical protein-protein interactions, relative to random expectation (Figure 1F–G). We therefore hypothesized that, in addition to its effects on ISG transcription, IFN stimulation induces widespread rewiring of protein-protein interaction networks.

### Quantitative interactome profiling of the type I IFN response

To map rearrangements in the human interactome induced by IFN stimulation, we applied a quantitative proteomic strategy based on protein correlation profiling (PCP) [24, 25] in combination with size exclusion chromatography (SEC). Under this workflow, protein complexes are separated by their size, and interacting protein pairs are inferred based on the similarity of their elution profiles (Figure 2A). The use of triplex SILAC labeling (SEC-PCP-SILAC) further enables differential analysis of protein-protein interactions between stimulated and unstimulated cellular states, to a high degree of quantitative precision [17].

**Figure 2.**
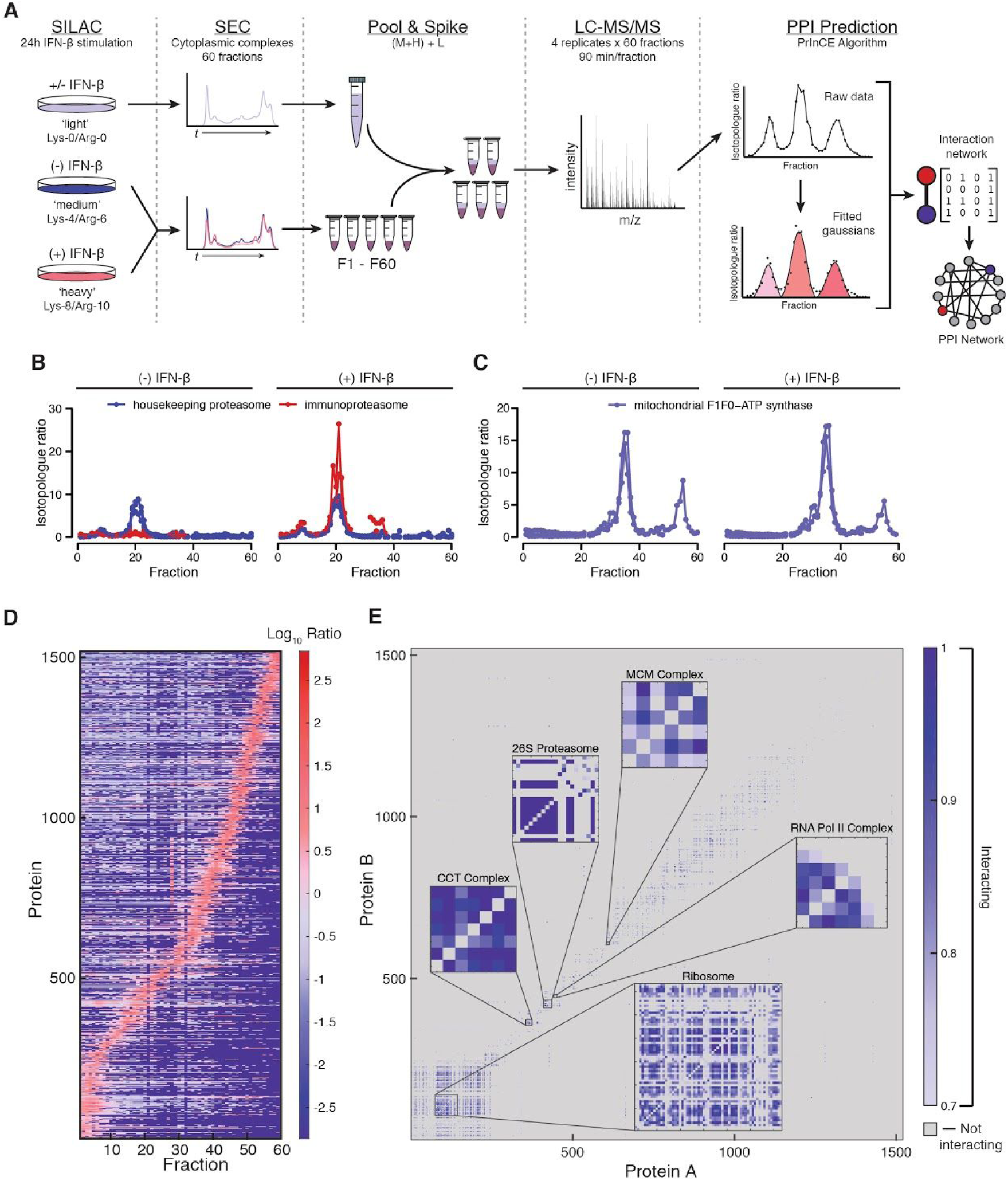
Mapping the interactome of IFN-stimulated cells by SEC-PCP-SILAC. (A) Schematic overview of the SEC-PCP-SILAC workflow. (B) PCP chromatograms of housekeeping proteasome and immunoproteasome-specific proteins in stimulated and unstimulated cells. (C) PCP chromatograms of mitochondrial F_1_F_0_–ATPase complex proteins in stimulated and unstimulated cells. (D) Complete set of PCP chromatograms (n = 1,520) defined by heavy/light ratio, arranged by index of maximum protein abundance, from a representative biological replicate. (E) Protein-protein interaction adjacency matrix for all 27,694 interactions with a precision greater than 70%, colored by interaction precision. Insets show adjacency matrices of known protein complexes from the CORUM database [82].

We applied SEC-PCP-SILAC to simultaneously compare the interactomes of cells stimulated with IFN for 24 h, labeled with heavy isotopes, and unstimulated cells, labeled with medium isotopes (Figure 2A). Fractions from the light channel, which included both stimulated and unstimulated cells in order to maximize proteome coverage, were pooled and spiked into all fractions as an internal standard. Sixty fractions were collected from each of three biological replicates, and were individually subjected to liquid chromatography–tandem mass spectrometry (LC-MS/MS) analysis. The resulting dataset was processed as a single experiment via MaxQuant [26] at a peptide and protein false discovery rate (FDR) of 1%, leading to the identification of 42,843 unique peptides from 2,590 protein groups across all 180 fractions (Figure 2D). Inspection of the PCP chromatograms revealed marked shifts between conditions for protein complexes with known roles in the innate immune response, such as the immunoproteasome (Figure 2B). Conversely, no differences between conditions were observed for housekeeping complexes such as the mitochondrial F_1_F_0_–ATP synthase, supporting the specificity of the technique (Figure 2C).

### Reconstruction of a high-confidence interactome network

To recover a high-confidence network of protein-protein interactions, we developed a multi-stage bioinformatic pipeline. First, whereas discussion of error rates in quantitative proteomics to date has focused primarily on errors in protein identification [27, 28], we observed a number of apparent errors in protein quantitation, some of which resulted in high-magnitude deviations in protein chromatograms (Figure S2C). Errors of this type have the potential to interfere with interaction detection, or to introduce spurious differential interactions between conditions. We therefore developed a network-based algorithm, ‘modern,’ to remove erroneous protein quantitations prior to further analysis (Methods). Application of modern to all three replicates led to removal of 1,308 erroneous protein quantitations (of 245,841 total quantitations, or 0.53%; Figures S2A and S2B).

Next, to derive a network of protein-protein interactions from the resulting chromatogram matrices, we applied PrInCE, a machine-learning pipeline for analysis of co-fractionation data [29, 30]. PrInCE fits a mixture of Gaussians to each chromatogram, then calculates a series of six features for each protein pair that reflect the likelihood of a physical interaction between those two proteins (Methods). These features are provided as input to a naive Bayes classifier, which calculates an interaction probability for every pair. Importantly, PrInCE assigns the likelihood of putative interactions based solely on the chromatograms themselves, without incorporating additional evidence from published functional genomics datasets, in contrast to several other approaches [31–33]. This approach facilitates unbiased detection of novel protein-protein interactions, without sacrificing discriminative power [34]. At a precision of 70%, PrInCE recovered a total of 27,694 binary protein interactions in IFN-stimulated and unstimulated cells (Figure 3A and Table S3). Visualization of the adjacency matrix revealed a sparse network of physical interactions, and confirmed recovery of well-known cellular protein complexes, such as the 26S proteasome, the chaperonin containing TCP-1 complex, and RNA polymerase II (Figure 2E).

**Figure 3.**
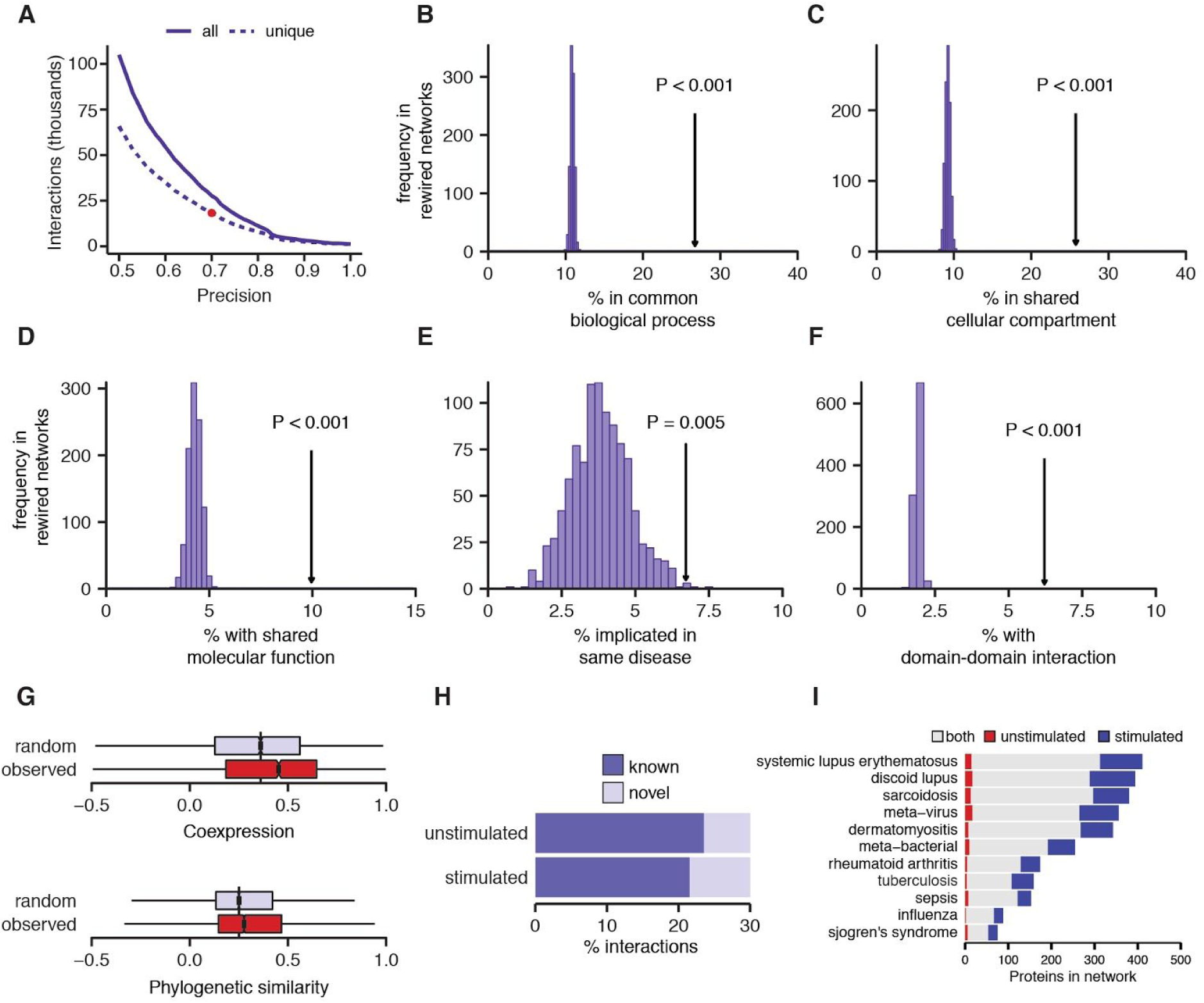
Biological relevance of the IFN interactome. (A) Precision-recall curve of all protein-protein interactions from unstimulated and IFN-stimulated networks (solid line), or unique interactions only (dashed line). (B–D) Proportion of interacting protein pairs sharing at least one biological process (B), cellular compartment (C), or molecular function (D) Gene Ontology term in the IFN interactome, arrow, or 1,000 randomly rewired networks, histogram. (E) Proportion of interacting protein pairs implicated in the same disease in the IFN interactome, arrow, or 1,000 randomly rewired networks, histogram. (F) Proportion of interacting protein pairs supported by a domain-domain interaction [35] in the IFN interactome, arrow, or 1,000 randomly rewired networks, histogram. (G) Pearson correlations reflecting protein abundance and phylogenetic profile similarity between interacting protein pairs in the IFN interactome or a randomly rewired network. (H) Proportion of previously known interactions in unstimulated and IFN-stimulated cells. (I) Number of genes found to be differentially expressed in meta-analyses of gene expression in eleven infectious or autoimmune diseases that were identified in the unstimulated or IFN-stimulated networks, or both.

### Biological relevance of the IFN interactome

We evaluated the overall biological relevance of the IFN-induced interactome by quantifying the degree to which interacting protein pairs tend to be involved in the same biological functions, localize to the same cellular compartments, or share the same molecular activities (Figure 3B–D). In all cases, we observed highly significant enrichments for shared GO terms between interacting pairs, relative to randomly rewired networks (all p < 0.001, permutation test). Further, we found interacting protein pairs were significantly more likely than random expectation to be implicated in the same disease (p = 0.005, permutation test; Figure 3E), and had more correlated patterns of protein abundance and phylogenetic profiles than non-interacting pairs (p < 10^−15^, Brunner–Munzel test; Figure 3G). Finally, interacting proteins were significantly more likely to share pairs of protein domains observed to physically interact in three-dimensional structural data [35], reflecting the power of SEC-PCP-SILAC to resolve physical protein-protein interactions, and not only functional associations (p < 0.001, permutation test; Figure 3F). Despite this enrichment, however, the majority of interactions detected were novel. Intriguingly, the IFN-stimulated network was significantly depleted for known interactions, relative to the unstimulated network (p = 8.0 × 10^−5^, χ^2^ test; Figure 3H), suggesting IFN stimulation specifically induces as-of-yet unmapped protein-protein interactions. Thus, multiple orthogonal lines of evidence support the high quality of our IFN interactome map, despite its recovery independent of any existing biological information.

Given that aberrant IFN signalling has been implicated in a broad range of infectious or autoimmune diseases, we further asked whether the IFN interactome could be used to interpret existing molecular datasets relevant to human pathologies. We drew on a resource of multi-cohort gene expression meta-analyses for 103 diseases [36, 37] to identify genes with reproducible evidence of differential expression in eleven diseases characterized by an elevated IFN transcriptional signature [8], including viral infections, systemic and discoid lupus erythematosus, rheumatoid arthritis, sarcoidosis, and Sjogren’s syndrome. We mapped protein-protein interactions for dozens to hundreds of differentially expressed genes from each disease (Figure 3I); notably, interactions for many such gene products were identified exclusively in the IFN-stimulated condition. Thus, the IFN interactome provides a reference to understand the consequences of dysregulated IFN signalling in a diverse range of human pathologies, by placing transcriptional markers of auto-inflammatory disease into a functional context.

### Evolutionary plasticity of the IFN response is mirrored at the interactome level

Comparative genomics approaches have highlighted genes involved in pathogen defense and the innate immune response as rapidly evolving, with divergence in both coding and regulatory sequences across species [38–40]. However, it remains unclear how this evolutionary divergence at the sequence level ultimately manifests at the interactome level. We quantified the degree to which each protein is ‘rewired’ upon IFN stimulation within the human interactome by calculating its autocorrelation between stimulated and unstimulated networks [41] (Methods). The tier of proteins with the lowest autocorrelation scores included several proteins with well-established roles in the innate immune response, such as the IFIT proteins and components of the immunoproteasome (Figure S3A). Comparing the IFN-induced autocorrelation of each protein to its evolutionary rate, as quantified by the ratio of nonsynonymous to synonymous substitutions (dN/dS), revealed a significant negative correlation (Spearman’s ρ = –0.11, p = 1.9 × 10^−4^; Figure S3B). Similar conclusions were reached by comparisons to the total number of species in which an ortholog of a given gene was present (ρ = 0.14, p = 5.3 × 10^−7^; Figure S3C) [42], or to the pLI score [43], a measure of mutational constraint derived from large-scale human exome sequencing (ρ = 0.18, p = 5.9 × 10^−5^; Figure S3D). Collectively, these results indicate that rapidly evolving proteins are disproportionately rewired in the protein-protein interaction network by IFN signalling, suggesting that species-specific differences in the innate immune response may be mediated in part through the effects of protein sequence divergence on protein-protein interactions.

### Functional landscape of IFN signalling

Despite the high quality of our IFN interactome, high-throughput maps of protein-protein interactions are unavoidably characterized by both false positives and false negatives [44]. We therefore sought to characterize the impact of IFN signalling on the human protein-protein interaction network more broadly, by developing a statistical framework for differential network analysis at the functional level (Figure S4A). Briefly, our approach first calculates the number, *n_PPI_*, of protein-protein interactions in both the stimulated and unstimulated networks involving proteins associated with a functional category of interest, and the difference between them, Δ*n_PPI_*. To assess statistical significance, the observed Δ*n_PPI_* is compared to a randomized distribution obtained from the *n_PPI_* values of 1,000 randomly rewired networks. The network rewiring procedure controls for biases stemming from network topology that may be independent of functionally relevant patterns [23, 45, 46].

A total of 407 GO terms were significantly enriched in either stimulated or unstimulated networks at 10% FDR, and were visualized as an enrichment map [47] (Figure 4 and Table S4). As expected, we observed a significant enrichment for interactions between proteins involved in the innate immune response in the IFN-stimulated network, including GO terms such as ‘antigen processing and presentation’ (Figure 4 and Figure S4B). The Wnt signalling pathway was likewise enriched in the IFN-stimulated network, consistent with its link to type I IFN signalling [48, 49] (Figure S4B). Intriguingly, the stimulated network was also enriched for terms relating to RNA processing and splicing, suggesting a role for physical interactions in the regulation of alternative splicing during the host response to viral infection [1, 50]. Conversely, the stimulated network was depleted for interactions involving chromatin remodelling and histone-modifying machinery, as well as interactions involved in negative regulation of ubiquitination. Surprisingly, we observed a significant enrichment for interactions related to translation and ribosome biogenesis upon IFN stimulation (Figure 4 and Figure S4B), potentially accounting for the apparent lag observed in our shotgun proteomics experiment between ISG transcription and translation. Collectively, these data reflect wide-ranging functional changes within the human interactome induced by IFN signalling.

**Figure 4.**
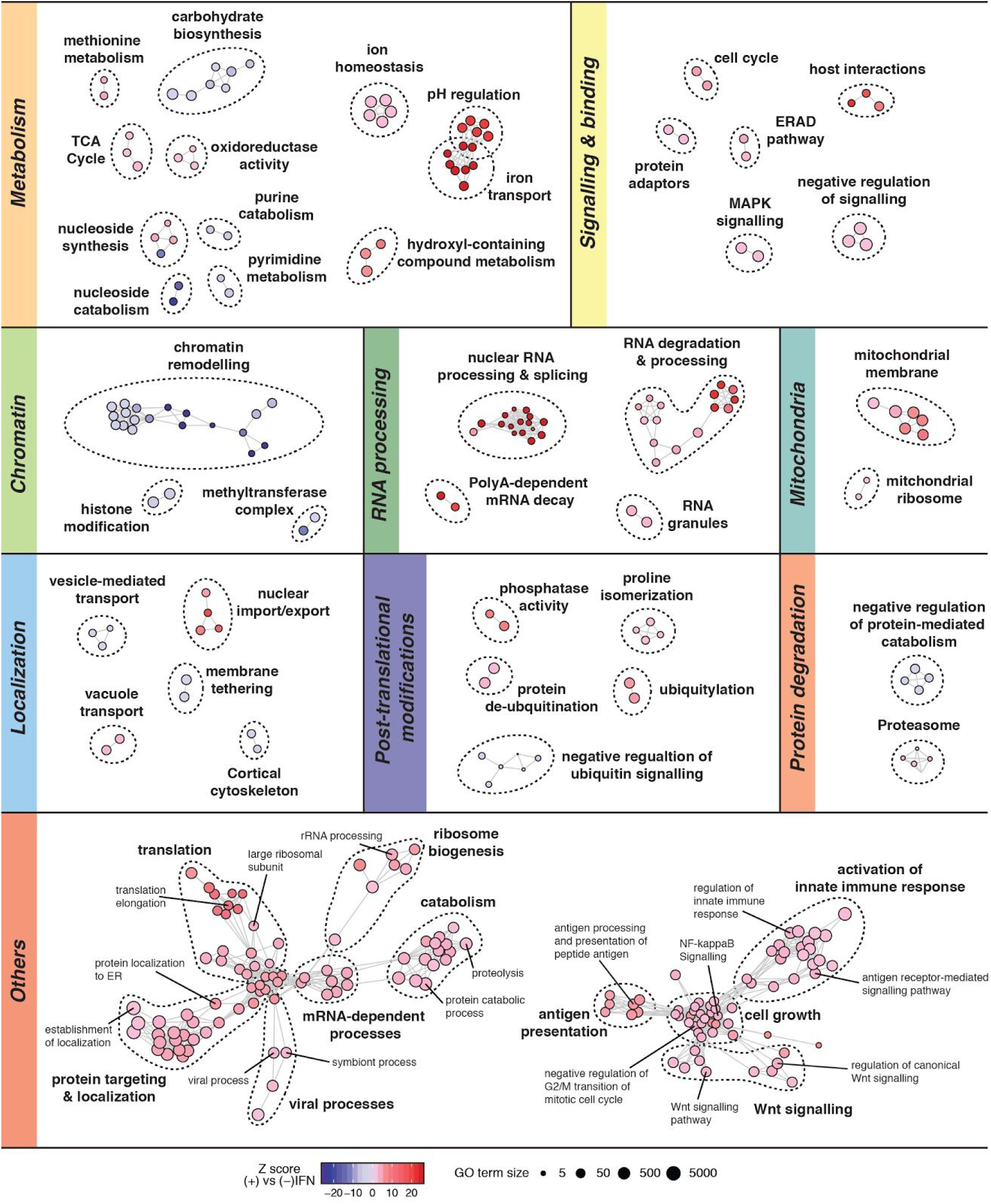
Differential network analysis of IFN-stimulated and unstimulated interactomes. Enrichment map of GO terms significantly enriched (red) or depleted (blue) in the IFN-stimulated interactome relative to the unstimulated interactome, at 10% FDR. Edges are defined between GO terms annotated to a common number of genes corresponding to a proteome-wide Jaccard index ≥ 0.33. Nodes clustered together in space therefore represent groups of functionally related GO terms, and are grouped by major biological processes or pathways.

### Interactions between evolutionarily conserved ISGs modulate the type I IFN response

Large-scale transcriptomic studies of the innate immune response across species and cell types have contrasted extensive heterogeneity in species- or cell type-specific transcriptional programs with a core module of universally upregulated genes, anchoring the transcriptional response across evolutionary and cellular contexts [7–9]. We sought to characterize the properties of evolutionarily conserved and species-specific ISGs at the interactome level. Using data from a comparative transcriptomic study of ten vertebrates [7], we defined sets of genes upregulated by IFN in all species (“core ISGs”), in human and at least one other species (“conserved ISGs”), or in humans only (“human-specific ISGs”). Strikingly, only the core set of evolutionarily conserved ISGs displayed a significant shift in chromatogram correlation upon IFN stimulation (p = 6.9 × 10^−3^, Brunner–Munzel test; Figure 5A), suggesting this evolutionarily ancient module of ISGs is induced to form physical interactions or protein complexes during the IFN response. To experimentally validate this result, we performed co-immunoprecipitations of four core ISGs in unstimulated or IFN-stimulated cells (Figure 5B). In three of four co-immunoprecipitations, core ISGs were significantly and selectively enriched among interactors induced by IFN stimulation, relative to conserved or human-specific ISGs (gene set enrichment analysis, p ≤ 5.1 × 10^−3^; Figure 5C).

**Figure 5.**
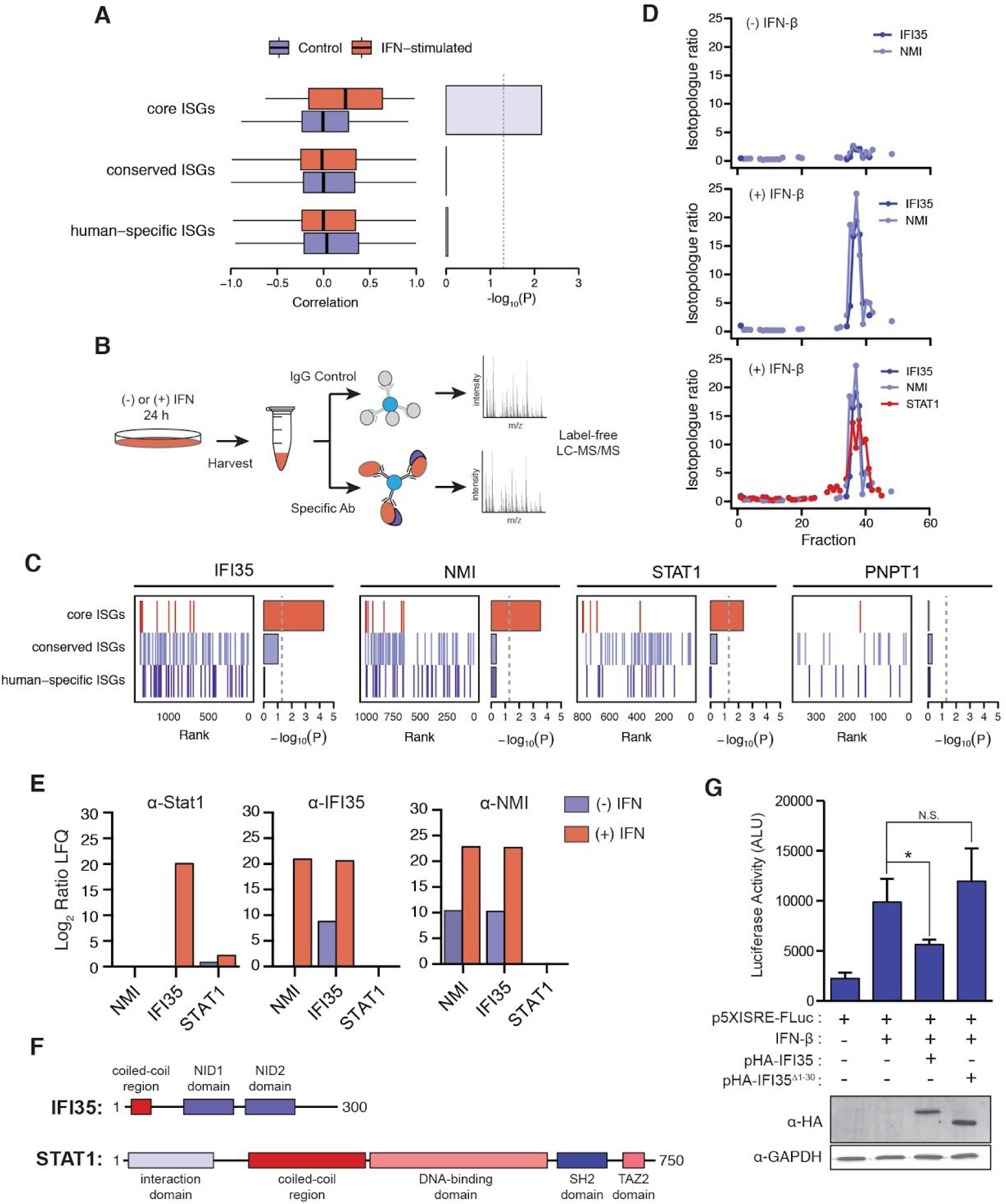
Interactions between evolutionarily conserved ISGs modulate the transcriptional response to IFN stimulation. (A) Left, distribution of correlations between pairs of proteins from core, conserved, and human-specific ISG sets in PCP chromatograms. Right, negative base-10 logarithm of p-values from Brunner–Munzel tests of the difference in medians. (B) Schematic overview of the affinity purification–mass spectrometry experiments of specific core ISGs in IFN-stimulated or unstimulated cells. (C) Gene set enrichment analysis barcode plots [113], showing ranks of core, conserved, and human-specific ISG products in comparisons of immunoprecipitations of IFI35 (left), NMI (middle), and STAT1 (right) from IFN-stimulated or unstimulated cells, alongside negative base-10 logarithms of p-values for each ISG set. (D) PCP chromatograms of IFI35, NMI, and STAT1 in IFN-stimulated and unstimulated cells. (E) Abundance of immunoprecipitated baits (STAT1, IFI35, or NMI), relative to an IgG control, in IFN-stimulated or unstimulated cells by label-free quantification (LFQ). (F) Protein domain content of IFI35 (top) and STAT1 (bottom). (G) Top, luciferase activities from cells transfected with a 5×ISRE-Fluc reporter construct to monitor STAT1 transcriptional activity (±S.D.). Cells were transfected with equal amounts of DNA in each case. Bottom, Western blots of HA-tagged IFI35 expression from lysates transfected with reporter constructs. * *P* < 0.05.

To shed light on the functional consequences of physical interactions between core ISGs, we focused on the interaction between IFI35 and STAT1, which was detected in IFN-stimulated cells by both SEC-PCP-SILAC (Figure 5D and Table S3) and co-immunoprecipitation of STAT1 (Figure 5E). Notably, we could not recover STAT1 in immunoprecipitations of IFI35, although we could recover its known binding partner NMI [51, 52]. Immunoprecipitations of NMI, which has previously been shown to interact with STAT1 during IFNγ treatment [53], likewise recovered IFI35, but not STAT1 (Figure 5E). These observations suggest that the interaction between STAT1 and IFI35 may be substoichiometric or potentially IFNβ-specific.

Given the role of IFI35 in repression of the IFN response by promoting degradation of RIG-I and decreasing IFNβ production [54, 55], we hypothesized that the interaction between IFI35 and STAT1 functioned to reduce STAT1 transcriptional activity as a potential mechanism to downregulate the IFN response. To investigate this hypothesis experimentally, we made use of a reporter construct containing 5× Interferon-<u>S</u>ensitive <u>R</u>esponse <u>E</u>lements (ISREs) fused to a firefly luciferase gene to monitor STAT1 transcriptional activity. Cells were transfected with the reporter construct, followed by transfection with a construct overexpressing hemagglutinin (HA)-tagged IFI35 and subsequent IFN stimulation. Control cells displayed low levels of luciferase activity that increased substantially upon IFN stimulation, as expected (Figure 5G). Expression of an N-terminal HA-tagged IFI35 resulted in a significant decrease in luciferase activity, suggesting that STAT1 transcription is impaired (Figure 5G). To further understand the mechanism underlying STAT1 transcriptional repression, we hypothesized that the IFI35–STAT1 interaction was mediated by the N-terminal coiled-coil domain of IFI35, with the internal coiled-coil domain of STAT1 its likely interaction partner. Consistent with this hypothesis, truncation of the coiled-coil domain from IFI35 recovered luciferase activity in transfected cells, despite expression of the truncated protein at similar levels as the full-length (Figure 5G). Taken together, these data suggest that IFI35 interacts with STAT1 at substoichiometric levels to fine-tune the transcriptional response to IFN stimulation.

### A specialized translational program mediated by RPL28 buffers ISG synthesis

Motivated by the observation of significant enrichment for interactions involved in translation and ribosome biogenesis in the IFN-stimulated interactome (Figure 4 and Figure S4B), we investigated the relationship between the mRNA and protein levels of ISGs in greater detail. Examining the expression of ISGs at the mRNA level in a densely sampled time-course transcriptomic experiment [8], we found proteins that were differentially expressed after 4 h of IFN stimulation reached peak transcriptional levels between 0.5–3.5 h (Figure S5A). Surprisingly, however, a large proportion of proteins differentially expressed at 24 h had peak mRNA expression at similar time points (Figure S5B), suggesting a lag in their translation. Based on these observations, we hypothesized that a specialized translational program, potentially mediated by changes in ribosome composition [56], is involved in establishing the IFN response.

To address this hypothesis, we performed sucrose density gradients to isolate free (40S, 60S, and 80S) and actively translating ribosomes (polysomes) in IFN-stimulated or unstimulated cells (Figure 6A). Polysome traces revealed an increase in free ribosomes upon IFN stimulation, apparent from the heightened 40S and 60S peaks, but no detectable changes in polysome levels (Figure 6B). To investigate changes in ribosome composition, fractions containing free ribosomes and polysomes, respectively, were pooled and subjected to quantitative mass spectrometry. Most ribosomal proteins were incorporated at similar levels in IFN-stimulated and unstimulated samples (Figure 6C and Table S5). However, we observed a substantial and selective increase in RPL28 incorporation upon IFN stimulation (Figure 6C). We confirmed this observation by Western blots of isolated ribosomes from IFN-stimulated or unstimulated cells. A substantial increase in RPL28 was observed compared to a control ribosomal protein, RPL14, suggesting RPL28 is selectively incorporated into ribosomes during the IFN response (Figure 6D).

**Figure 6.**
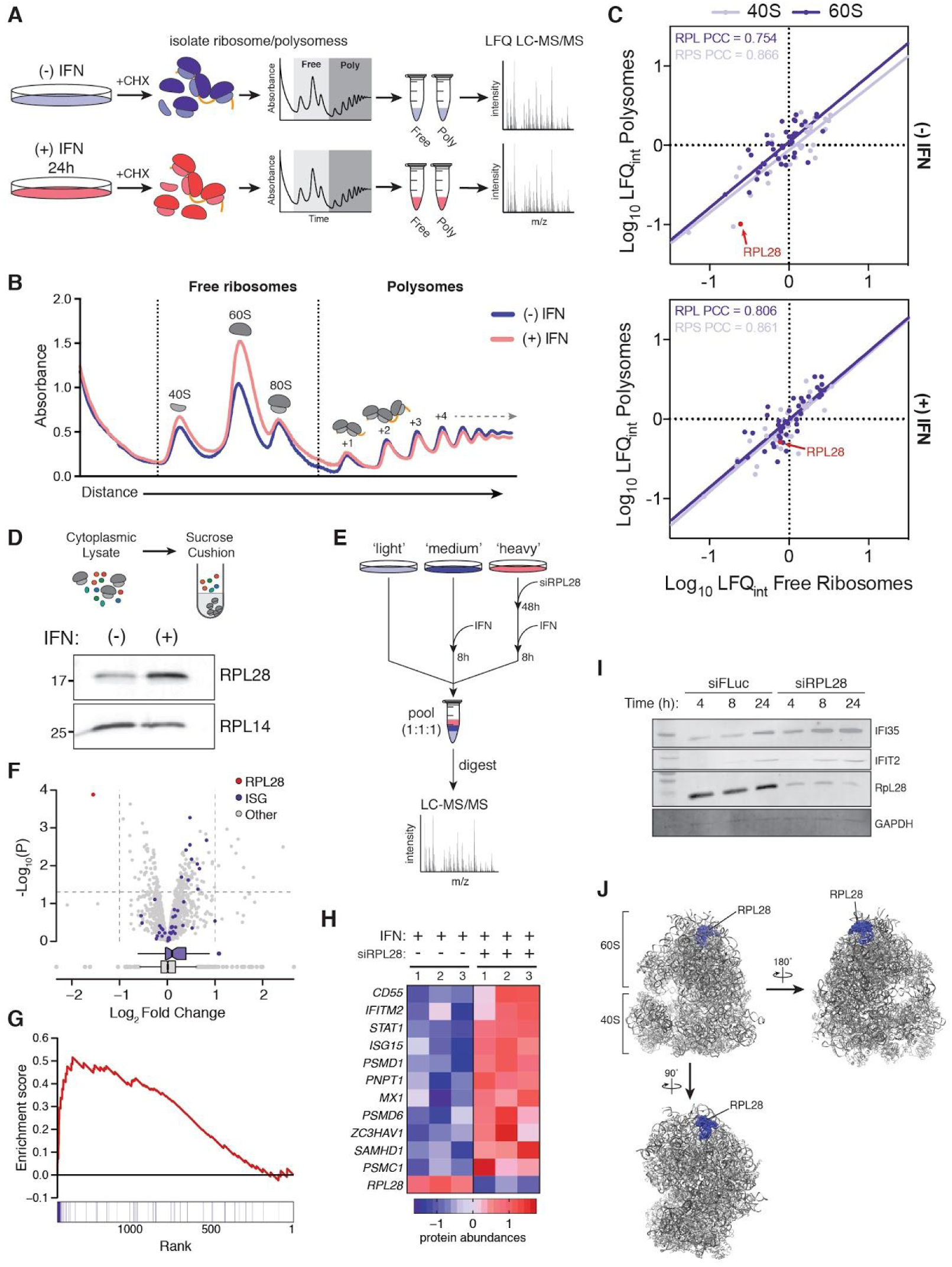
IFN stimulation induces changes in ribosome composition to regulate ISG synthesis. (A) Schematic overview of the sucrose gradient experiments to determine changes in composition of free (40S, 60S, and 80S) and actively translating ribosomes (polysomes). (B) Representative traces of sucrose density gradients from cells stimulated with IFN for 24 h or unstimulated cells. (C) Median protein abundance across three replicates from pooled free ribosome or polysome fractions by label-free quantification (LFQ) in IFN-stimulated or unstimulated cells. PCC, Pearson correlation coefficient. (D) Western blot of ribosomes isolated from IFN-stimulated or unstimulated cells by sucrose cushion. (E) Experimental workflow for shotgun proteomics analysis on RPL28-depleted cells after IFN-stimulation. Yea (F) Volcano plot showing differential protein abundance in cells stimulated with IFN for 8 h and treated with siRPL28, relative to controls. (G) Gene set enrichment analysis enrichment score, top, and barcode plot [113], bottom, showing ranks of ISGs in a comparison siRPL28-treated and IFN-stimulated cells compared to untreated controls. (H) Heatmap showing abundance of select well-studied ISGs in siRPL28-treated and IFN-stimulated cells compared to untreated controls. (I) Western blots of select ISGs in cells treated with siRPL28 or control siRNA after 4 h, 8 h, and 24 h of IFN stimulation. (J) Crystal structure of the human 80S ribosome (PDB: 4UG0), with RPL28 highlighted in blue.

We next sought to characterize the functional consequences of RPL28 incorporation during the IFN response. Over the past decade, it has become clear that ribosomes do not exist as a single homogenous population within an organism, but instead undergo dynamic changes in composition, with incorporation of individual ribosomal components forming ‘specialized ribosomes’ that promote the translation of specific mRNA classes [57–60]. Based on this body of evidence, and the fact that RPL28 lies in a solvent-accessible portion of the ribosome (Figure 6J), we hypothesized that IFN-dependent RPL28 incorporation facilitates translation of mRNAs related to the type I IFN response. In support of this hypothesis, examination of the Human Protein Atlas [61] and FANTOM5 database [62] indicated RPL28 is expressed at the highest levels in human immune system tissues, such as the thymus and lymph nodes (Figure S5C–D), consistent with an immunological role.

To determine the functional role of RPL28, we used siRNAs to knockdown RPL28 followed in the context of IFN stimulation to precisely quantify changes in protein synthesis. Cells labeled with heavy isotopes were treated with siRPL28 for 48 h, after which both medium- and heavy-labeled cells were exposed to IFN for 8 h, with unstimulated light-labeled cells serving as a baseline control (Figure 6E). A total of 1,940 proteins were identified, of which 1,421 were quantified in at least two of three replicates (Table S6). To our surprise, comparison of siRPL28-treated cells to controls revealed an increase in ISG abundance upon RPL28 knockdown (Figures 6F and 6H), an effect which was highly significant by gene set enrichment analysis (p = 4.0 × 10^−4^; Figure 6G). We further confirmed this finding by Western blot, observing increases in ISG abundance over time in the RPL28 knockdown compared to an siRNA control (Figure 6I). Importantly, to rule out the possibility that RPL28 leads to impaired ribosome biogenesis and global downregulation of translation rates (Mills and Green, 2017), we monitored protein synthesis by metabolic labeling with [35S]-Met/Cys, finding that RPL28 depletion did not alter global protein levels in either our SILAC experiment (Figure 6F) or when assessed with metabolic labeling (Figure S5E), suggesting that the ribosome remains functionally competent. It is possible that the changes observed in ISG abundance are merely due to an increase in mRNA levels. To further rule out the possibility that the changes in ISG protein abundance are mediated primarily by an increase in mRNA levels, we performed qRT-PCR, which did not reveal a significant difference between siRPL28- or control-treated cells (with the exception of *NMI*; Figure S5F). Taken together, these results suggest that RPL28 is specifically incorporated into ribosomes upon IFN stimulation, where it acts to selectively downregulate ISG synthesis.

## DISCUSSION

The pleiotropic effects of type I IFN stimulation on transcriptional regulation have been appreciated for over two decades [2]. In turn, the maturation of increasingly sensitive technologies for transcriptome profiling, complemented by functional assays, have led to the identification of hundreds of ISGs [6]. Yet, with relatively few exceptions, the functional roles of these effectors in the antiviral response remain incompletely delineated. In particular, little is known about how existing cellular networks are influenced by IFN stimulation, and how newly synthesized ISGs engage these complex networks. Here, we have used protein correlation profiling to construct a differential network map of the human protein-protein interactome in response to type I IFN signalling. This network map identifies specific interactions for known ISGs under homeostatic conditions, and reveals patterns of interaction rewiring induced by IFN stimulation. The IFN interactome thus places ISGs into a functional context, providing a platform for further mechanistic dissection of their roles in the innate immune response. As one example, we find that IFI35 binds to STAT1 to fine tune its transcriptional activity, likely through a coiled-coil interaction, implicating IFI35 as a direct negative regulator of the IFN response. Moreover, given the dysregulation of IFN signalling in both monogenic diseases [5], as well as a broader spectrum of autoimmune and neuropsychiatric disorders [4], our work provides a framework to develop a deeper understanding of the mechanisms that protect against inappropriate immune activation.

Charting macromolecular interaction networks in a physiologically relevant and differential context, particularly at the proteome scale, represents a longstanding challenge [16]. Our results highlight the unique power of protein correlation profiling, in combination with SILAC labelling and size exclusion chromatography, to systematically resolve interaction dynamics in the innate immune response, or in response to cellular perturbations more broadly [17, 63]. Widely used methods for interactome mapping, such as yeast two-hybrid or affinity purification-mass spectrometry, rely on heterologous expression of fusion proteins or the introduction of a protein tag; the former removes proteins completely from their endogenous cellular context, whereas the latter can disrupt the native interactions or subcellular localization of the tagged protein [64]. Thermal proximity co-aggregation (TPCA) has demonstrated promise for interrogating protein interaction networks *in vivo*, or across distinct cellular states [65–67], but to date has been limited to monitoring the dynamics of known interactions or protein complexes, rather than enabling *de novo* network inference. In contrast, the primary disadvantage associated with SEC-PCP-SILAC is its moderate bias towards proteins of greater cellular abundance [68]. In combination with the computational tools for differential network analysis described here, SEC-PCP-SILAC represents a powerful, untargeted approach to define rearrangements in the human interactome in response to cellular stimuli.

Our functional analysis of the IFN-induced differential interactome implicated rewiring of interactions involved in protein translation and ribosome biogenesis in the IFN signalling cascade, leading us to uncover a novel regulatory mechanism involved in the innate immune response. We find that increased incorporation of RPL28 into the ribosome upon IFN stimulation represses synthesis of ISGs, whereas global translation remains unaffected. Given the need for tight regulation of the IFN response to avoid aberrant over-activation, RPL28 may act as a buffer on excessive ISG translation, in effect imposing a secondary layer of regulation beyond mRNA transcription. Overall, this finding adds to a growing body of literature demonstrating how modulation of ribosome composition can facilitate translational control of specific mRNA classes [58–60].

## CONCLUSIONS

In sum, the map of the IFN-induced interactome presented here systematically expands our understanding on the organization of the innate immune response while complementing previous functional and systems-level studies, providing a rich resource to inform hypothesis-driven experiments. Our data reveals a surprisingly broad spectrum of rewiring in cellular pathways induced by IFN, and uncovers novel regulatory mechanisms at the levels of transcription and translation that modulate the IFN response. Intersecting this network map with data from genome-wide association or exome sequencing studies could prove an effective strategy to further understand how these pathways are perturbed by common or rare variants in the context of autoimmune or neuropsychiatric diseases [69, 70]. More broadly, our work establishes a proteomic and bioinformatic platform to delineate the complex networks of regulatory pathways activated in response to physiological or pathophysiological stimuli.

## DECLARATIONS

### AVAILABILITY OF DATA

The mass spectrometry proteomics data have been deposited to the ProteomeXchange Consortium [110] via the PRIDE partner repository [111] with the dataset identifier PXD013809. In addition, processed chromatograms for each replicate have been deposited to the EMBL–EBI BioStudies database [112], with accession S-BSST254.

### AUTHOR CONTRIBUTIONS

Conceptualization, C.H.K, M.A.S. and L.J.F.; Methodology, C.H.K., M.A.S. and L.J.F.; Software, M.A.S. and R.G.S.; Validation, C.H.K., M.A.S., A.M.M., D.D.T.A.; Formal Analysis, M.A.S. and C.H.K.; Investigation, C.H.K., M.A.S., A.M.M, D.D.T.A., and Q.W.T.C.; Resources, E.J. and L.J.F.; Writing - Original Draft, C.H.K., M.A.S., and L.J.F.; Writing, Reviewing & Editing - C.H.K., M.A.S., A.M.M, D.D.T.A., Q.W.T.C., N.S., R.G.S., E.J., and L.J.F.; Supervision, E.J. and L.J.F.; Funding Acquisition, E.J. and L.J.F.

### FUNDING

This work was supported by Canadian Institutes of Health Research (CIHR) grant and National Sciences and Engineering Research Council (NSERC) Discovery grants awarded to L.J.F. and E.J. C.H.K. was supported by an NSERC Post-Graduate Scholarship and a UBC Four Year Fellowship. M.A.S. is supported by a CIHR Vanier Canada Graduate Scholarship, an Izaak Walton Killam Memorial Pre-Doctoral Fellowship, a UBC Four Year Fellowship, and a Vancouver Coastal Health–CIHR–UBC MD/PhD Studentship. Mass spectrometry infrastructure used here was supported by Genome Canada/Genome BC (214PRO).

## ACKNOWLEDGEMENTS

We thank Nichollas Scott (University of Melbourne) for advice on experimental design and assistance in shotgun proteomics data acquisition and Kyung-Mee Moon for assisting with manuscript preparation. We thank Dr. Curt Horvath (Northwestern University) for the 5×ISRE-Fluc reporter construct. We thank the members of the Foster lab for invaluable discussions.

## COMPETING INTERESTS

The authors have no competing interests.

## METHODS

### Cell culture and IFNβ stimulation

HeLa cells were cultured in Dulbecco’s modified Eagle medium (DMEM) supplemented with 10% fetal bovine serum (FBS), 1× penicillin-streptomycin (Pen-Strep), and 2 mM L-Glutamine at 37°C. For IFNβ stimulation, cells were seeded and incubated overnight at 37°C. The following day, cells were washed with 1× Phosphate buffered saline (PBS) and stimulated with 1000 U/mL of human recombinant IFNβ (R&D Systems) in DMEM for the designated length of time. For SILAC experiments, cells were stimulated in the appropriate SILAC-formulated media.

### Plasmids and transfections

HA-tagged IFI35 was generated as follows. First, total RNA was isolated from IFNβ-stimulated (24 h) HeLa cells via TRIzol extraction (Thermo Fisher) followed by RT-PCR using an oligo dT primer. Desired sequences were amplified from cDNA using primers IFI35-F (5’ – TAGGGTACCATGTCAGCCCCACTGGATGCCG – 3’), and IFI35-R (5’ – TAGCTCGAGCTAGCCTGACTCAGAGGTGAAGACTGC – 3’). Amplicons were digested, followed by ligation into pcDNA3.1. Subsequently, a 3×HA tag was N-terminally fused onto STAT1 and IFI35, respectively, by amplifying the cloned sequences with primers that incorporated the tags and sub-cloning into the respective constructs. Constructs were confirmed through sequencing. The ISRE-Luc reporter construct was a gracious gift from Dr. Curt Horvath (Northwestern University) and contains 5× ISG54 ISRE elements upstream of a TATA box and a firefly luciferase open reading frame.

Transfections were done as follows: briefly, 3.0 × 10^5^ HeLa cells were seeded into 6-well plate and incubated for 24 h at 37°C. Plasmids (2 µg) and transfection reagent (5 µL of Lipofectamine 2000; Invitrogen) was added to 125 µL of OptiMEM serum-free media (ThermoFisher) in separate tubes and incubated for 5 min. Tubes were combined and incubated for 15 min. Media was aspirated from cells and the 250 µL transfection mix was added to cells dropwise. Complete media was added to each well and cells were incubated at 37°C. For RPL28 and control siRNA experiments, cells were transfected as per manufacturer’s’ protocol at a final concentration of 25 nM using Dharmafect I transfection reagents (Dharmacon).

### Luciferase assays

Luciferase assay were carried out using a Luciferase Assay System Kit (Promega). Briefly, cells transfected in a 6-well plate as described above were harvested and washed with 1× PBS. Cells were then lysed using 1× Passive Lysis Buffer as per manufacturer’s protocol (Promega). Protein concentration was then determined via Bradford assay. Equal amounts of protein (30 µg) was added to a Costar Flat White 96-well plate for each condition and samples were brought to equal volume with 1× Passive Lysis Buffer. Following this, 50 µL of luciferase reagent was added to each well and luminescence was recorded on an Infinite M200 microplate reader (Tecan).

### Western blots

Equal amounts of protein were resolved on a 12% SDS-PAGE gel and then transferred to a polyvinylidene difluoride Immobilon-FL membrane (PVDF; Millipore). Membranes were blocked for 30 min at room temperature with 5% skim milk in TBST (50 mM Tris, 150 mM NaCl, 1% Tween-20, pH 7.4). Blots were incubated for 24 h at 4°C with the following antibodies: mouse anti-GAPDH (1:1000; AbLab), rabbit anti-HA (1:1000; Cell Signalling - C29F4), rabbit anti-RPL14 (1:1000; Bethyl Laboratories - A305-052A), rabbit anti-RPL28 (1:1000; AbCam - ab138125), mouse anti-IFIT2 (1:1000; Santa Cruz - sc-390724), mouse anti-IFI35 (1:1000; Santa Cruz - sc-100769), or rabbit anti-NMI (1:1000; AbCam - ab183724). Membranes were washed 3 times with TBST and incubated with either IRDye 800CW goat anti-mouse (1:5000; Li-Cor Biosciences), or IRDye 800CW goat anti-rabbit (1:5000; Li-Cor Biosciences) for 1 h at room temperature. Membranes were then washed 3 more times with TBST before imaging on an Odyssey imager (Li-Cor Biosciences).

### Metabolic labeling

HeLa cells were transfected with siFluc or siRPL28 for 48 h before being stimulated with IFNβ for 4 or 8 h and labelled with 250µCi [^35^S]-Met/Cys for 30 min. Cells were washed twice with 1 mL PBS and harvested with 100 µL RIPA buffer. Equal amounts of lysates were loaded on 12% SDS-PAGE gels. Gels were dried and radioactive bands were analyzed using a phosphorimager (GE Amersham Typhoon). To quantify incorporated [^35^S]-Met/Cys, 20 µg of protein was precipitated with 25% trichloroacetic acid (TCA) before being filtered through a glass fibre filter. Subsequently, the filter was washed three times with 5% TCA followed by 100% acetone. The filter was suspended in scintillation fluid and analyzed on a liquid scintillation counter (Perkin Elmer).

### Polysome and ribosome isolation

HeLa cells (1.0 × 10^7^) were seeded in 150 mM tissue culture plates for each condition and incubated for 24 h at 37°C. Media was aspirated and replaced with either control media or media containing 1000 U/mL IFNβ and cells were incubated for 24 h at 37°C before being subjected to polysome or ribosome isolation. For whole ribosome isolation, cells were lysed with Ribosome Lysis Buffer (300 mM NaCl, 15 mM Tris-HCl, 6 mM MgCl_2_, 1% Triton X-100, 1 mg/mL Heparin, pH 7.5). Lysates were clarified by centrifugation at 20,000 r.c.f. for 10 min at 4°C. Clarified lysates were layered over a sucrose cushion at a 1:1 (v/v) ratio (2M Sucrose in Ribosome lysis buffer) followed by centrifugation at 100,000 r.c.f. for 24 h at 4°C. Pelleted ribosomes were resuspended in RIPA buffer (50 mM Tris, 150 mM NaCl, 0.1% SDS, 0.5% sodium deoxycholate, 1% Triton X-100, 0.5 mM EDTA) and protein concentration was quantified by Bradford assay (BioRad). Equal amounts of protein were used for Western blot analysis.

For polysome analysis, after treatment with IFNβ, cycloheximide (100 µg/mL) was added to the media and cells were incubated for 5 min. Cells were washed 3 times with 1× PBS plus cycloheximide (100 µg/mL) before being lysed in 400µL of Polysome Lysis Buffer (300 mM NaCl, 15 mM Tris-HCl, 15 mM MgCl_2_, 100 µg/mL cycloheximide, 1 mg/mL heparin). Lysates were clarified by serial centrifugation at 800 r.c.f. for 5 min at 4°C, then 13,000 r.c.f. for 10 min at 4°C. RNA was quantified via NanoDrop and equal amounts of RNA (500 µg) were loaded onto a linear 10-50% sucrose gradient made in Polysome Lysis Buffer. Gradients were centrifuged in a SW41 Ti Rotor (Beckman) for 2.5 h at 40,000 RPM at 4°C. Fractions were collected on a Gradient Station IP Fractionator (BioComp). Fractions (750 µL) corresponding to free ribosomes (40S, 60S, and 80S monosomes) and polysomes were pooled and proteins were precipitated using trichloroacetic acid. Resulting protein pellets were reduced, alkylated, and subjected to trypsin digestion before being analyzed by LC-MS/MS.

### SILAC labeling and shotgun mass spectrometry

HeLa cells were cultured in DMEM (Lys/Arg^-/-^) supplemented with 10% dialyzed FBS (Invitrogen), 1× Pen-Strep, and combinations of the following lysine and arginine isotopologues: for “light” (“L”)–labeled cells L-arginine (84 mg/L) and L-lysine (146 mg/L)(Sigma-Aldrich); for “medium” (“M”)-labeled cells, ^13^C_6_-L-arginine (87 mg/L) and D_4_-L-Lysine (150 mg/L); for “heavy” (“H”)-labeled cells ^13^C_6_^15^N_4_-L-arginine (89 mg/L) and ^13^C_6_^15^N_2_-L-lysine (154 mg/L)(Cambridge Isotope Laboratories). Cells were split into each SILAC formulation and passaged six times to allow for complete incorporation of amino acid isotopologues.

For shotgun proteomic analysis, ∼1.0 × 10^7^ cells were harvested from control (light), 4 h IFNβ stimulation (medium), and 24 h IFNβ stimulation (heavy). Cells were lysed in Lysis Buffer (4% SDS, 10 mM DTT, 100 mM Tris-HCl, pH 8.8) and heated at 95°C for 5 min. Samples were then centrifuged for 10 min at 16,000 r.c.f. at 4°C and the supernatant was collected. Protein concentrations were then measured via BCA assay (Thermo Fisher). 100 µg of protein from each sample (light, medium, and heavy) were combined and subjected to acetone precipitation. Protein pellets were resuspended in a 6 M/2 M urea/thiourea mixture. Samples were reduced and alkylated by adding 6 µg of DTT and 15 µg of iodoacetamide and incubating at room temperature in the dark for 30 min and 20 min, respectively. 3 µg of LysC was added to each sample and incubated for 3 h at room temperature. Subsequently, samples were diluted with 4 volumes of Digestion Buffer (50 mM NH_4_HCO_3_) and trypsin (Promega) was added at a ratio of 1:50. Samples were incubated shaking overnight at room temperature. The resulting peptide supernatant was acidified to pH < 2.5 and purified using homemade Stop-and-go-extraction tips (StageTips) composed of C18 Empore material (3 M) packed in to 200 µL pipette tips [71]. StageTips were conditioned with methanol and equilibrated with 1% trifluoroacetic acid (TFA; loading buffer). Peptide supernatants were loaded onto the columns and washed with two bed volumes of Buffer A (0.5% formic acid). Peptides were eluted with Buffer B (80% MeCN, 0.5% formic acid), dried down. Peptides from each biological replicate were then subjected to high pH reverse-phase (RP) fractionation on an Agilent 1100 HPLC system with an Agilent Zorbax Extend column (1.0×50 mm, 3.5 µm particles, flow rate of 50 µL/min). Dried peptides were resuspended in RP Buffer A (5 mM NH_4_HCO_2_, 2% MeCN, pH 10), injected, and eluted from the column over a 60 min gradient: 0 to 5 min 6% RP Buffer B (5 mM NH_4_HCO_2_, 90% MeCN); 5-7 min 8% RP Buffer B; 7-45 min 27% RP Buffer B; 45-49 min 31% RP Buffer B; 49-53 min 39% RP Buffer B; 53-60 min 60% RP Buffer B. The column was washed by running 100% RP Buffer B for 5 min. Fractions were collected every 40s for 60 min. Every eighth fraction was then concatenated, dried, and resuspended in Buffer A for mass spectrometry analysis.

Purified peptides were analyzed using an Easy nano LC 1000 nanoflow HPLC (Thermo Fisher) on-line coupled to a Q-Exactive mass spectrometer (Thermo Fisher). The LC was operated in a trapping mode (two column system) using a 4 cm long, 100 μm inner diameter fused silica trap column. The analytical column was from 75 μm inner diameter fused silica capillary and it was either with an integrated spray tip, or it was fritted and attached to a 20 μm inner diameter fused silica gold coated spray tip. Columns with spray tip and spray tips for fritted columns were pulled on a P-2000 laser puller from Sutter Instruments to 6 μm diameter opening. Added spray tips were coated on EM SCD005 Super Cool Sputtering Device (Leica). The trap column was packed with 5 μm diameter Aqua C-18 beads (Phenomenex) to 2 cm, while the analytical column was packed with 3.0 μm diameter Reprosil-Pur C-18-AQ beads (Dr. Maisch). The trap column was conditioned with 20 μL Buffer A and the analytical column was conditioned with 4 μL of the same buffer. Samples were loaded with 20 μL of Buffer A. The analysis was performed at 250 nL/min over 180 min with a gradient from 0% to 40% buffer B over 180 min, then from 40% to 100% over 2 min and held at 100% B over 10 min. The LC autosampler thermostat was set at 7°C. The Q-Exactive was operated in a data dependent mode using Xcalibur v.2.2 (Thermo Fisher) and set to acquire a full-range scan at 70,000 resolution from 350 to 2000 Th (AGC target 3E6) and to fragment the top ten multiply charged ions above 5% underfill ratio by HCD (resolution 17500, AGC target 1E5, maximum injection time 60 ms, NCE 28) in each cycle. Parent ions were then excluded from MS/MS for the next 25 s. Error of mass measurement is typically within 5 ppm and was not allowed to exceed 10 ppm.

### SEC-PCP-SILAC sample preparation

Cell lysis and size exclusion chromatography were performed as previously described [17, 63], with minor modifications. Briefly, after 24 h treatment with IFNβ cells were immediately harvested by centrifugation at 200 r.c.f. for 5 min at 4°C and washed three times with ice-cold 1× PBS. Cells of the same SILAC label were pooled and resuspended in 3 mL of ice-cold size-exclusion chromatography (SEC) buffer [50 mM KCl, 50 mM NaCH_3_COO, 50 mM Tris, pH 7.2, containing 1× EDTA-free HALT protease & phosphatase inhibitor cocktail (Thermo Fisher)]. Cells were lysed via Dounce homogenization for 2.5 min and insoluble material was removed by ultracentrifugation at 100,000 r.c.f. for 15 min at 4°C. Subsequently, the supernatants were concentrated over a 100 kDa molecular weight cutoff spin column (Sartoris Stedim, Goettingen, Germany). Equal amounts of protein from heavy-labeled and medium-labeled lysates were combined and immediately injected into a chromatography systems with two 300 × 7.8 mm BioSep4000 Columns (Phenomenex) equilibrated with SEC buffer. Samples were collected into 80 fractions by a 1200 Series analytical HPLC (Agilent Technologies) at a flow rate of 0.5 mL/min at 8°C. To avoid aggregated proteins and monomers, only fractions 6-65 were submitted for LC-MS/MS analysis and utilized for PCP. The light-labeled SILAC lysates consisted of unstimulated and IFN-stimulated cells to serve as an internal standard. These samples were independently separated by SEC from the medium/heavy samples. To generate the light reference mixture, fractions 6-65 were pooled and spiked equally into each of the corresponding medium/heavy fractions at a volume of 1:0.75 (medium/heavy to light). A urea/thiourea mix was added to protein fractions to create a final concentration of 6 M/2 M urea/thiourea. Samples were reduced and alkylated by adding 6 µg of DTT and 15 µg of iodoacetamide and incubating at room temperature in the dark for 30 min and 20 min, respectively. 3 µg of LysC was added to each sample and incubated for 3 h at room temperature. Subsequently, samples were diluted with 4 volumes of Digestion Buffer (50 mM NH_4_HCO_3_) and trypsin (Promega) was added at a ratio of 1:50. Samples were incubated shaking overnight at room temperature. The resulting peptides were purified via STAGE tips.

### Immunoprecipitation mass spectrometry sample preparation

HeLa cell lysates subjected to IP-MS analysis were prepared similarly as described for SEC-PCP-SILAC, apart from isotopologue labeling. In short, control and IFN-stimulated HeLa were harvested via centrifugation at 200 r.c.f. for 5 min at 4°C then washed three times with ice-cold PBS. Cells were then lysed via Dounce homogenization in SEC buffer followed by centrifugation at 100,000 r.c.f. Supernatants were concentrated over a 100 kDa molecular weight cutoff spin column (Sartoris Stedim, Goettingen, Germany). Protein concentration was determined by NanoDrop (Thermo Fisher). 250 µg of protein from control or IFN-stimulated samples was diluted to 500 µL in SEC buffer and incubated with the desired antibody or IgG as a control overnight at 4°C. Antibody concentrations were used as follows: mouse anti-IFI35 (1:50; Santa Cruz - sc-100769), rabbit anti-NMI (1:50; AbCam - ab183724), mouse anti-STAT1 (1:50; AbCam - ab3987), and mouse anti-PNPT1 (2 µg; Santa Cruz - ab-271479). Protein A/G Magnetic beads (25 µL; Thermo Fisher) pre-washed with SEC buffer were added to the samples and incubated 1 h at room temperature. Beads were washed three times with 20× bed volume of SEC buffer. Samples were then subjected to an on-bead in-solution trypsin digestion. Peptides were purified by STAGE tips.

### Tandem liquid chromatography mass spectrometry of SEC-PCP-SILAC and LFQ samples

Purified peptides were analyzed using a quadrupole time of flight mass spectrometer (Impact II; Bruker Daltonics) on-line coupled to an Easy nano LC 1000 HPLC (Thermo Fisher) using nanoBooster with methanol and a Captive spray nanospray ionization source (Bruker Daltonics) including a 2 cm long, 100 μm inner diameter fused silica fritted trap column, and 40 cm long, 75 μm inner diameter fused silica analytical column with an integrated spray tip (6-8 μm diameter opening, pulled on a P-2000 laser puller from Sutter Instruments). The trap column was packed with 5 μm Aqua C-18 beads (Phenomenex) while the analytical column was packed with 1.9 μm diameter Reprosil-Pur C-18-AQ beads (Dr. Maisch). Buffer A consisted of 0.1% aqueous formic acid, and buffer B consisted of 0.1% formic acid in acetonitrile. Samples were resuspended in buffer A and loaded with the same buffer. Standard 90 min gradients were from 0% B to 35% B over 90 min, then to 100% B over 2 min, held at 100% B for 15 min. Before each run, the trap column was conditioned with 20 μL buffer A, the analytical with 4 μL of the same buffer, and the sample loading was set at 20 μL (for samples up to 13 μL volume). The LC thermostat temperature was set at 7°C. The Captive Spray Tip holder was modified similarly to an already described procedure [72]. The fused silica spray capillary was removed (together with the tubing which holds it) to reduce the dead volume, and the analytical column tip was fitted in the Bruker spray tip holder using a piece of 1/16 in × 0.015 PEEK tubing (IDEX), an 1/16 in metal two-way connector and a 16-004 Vespel ferrule. The sample was loaded on the trap column at 850 Bar and the analysis was performed at 0.25 μL/min flow rate. The Impact II was set to acquire in a data-dependent auto-MS/MS mode with inactive focus fragmenting the 20 most abundant ions (one at a time at 18 Hz rate) after each full-range scan from m/z 200 Th to m/z 2000 Th (at 5 Hz rate). The isolation window for MS/MS was 2 to 3 Th depending on parent ion mass to charge ratio and the collision energy ranged from 23 to 65 eV depending on ion mass and charge [72]. Parent ions were then excluded from MS/MS for the next 0.4 min and reconsidered if their intensity increased more than 5 times. Singly charged ions were excluded since in ESI mode peptides usually carry multiple charges. Strict active exclusion was applied. Error of mass measurement is typically within 5 ppm and was not allowed to exceed 10 ppm. The nano ESI source was operated at 1900 V capillary voltage, 0.20 Bar nanoBooster pressure, 3 L/min drying gas and 150°C drying temperature. The cross connector between the trap column, waste out capillary and analytical column was grounded via a 0.4 mm platinum wire to prevent electrical corrosion of the LC S valve.

### Protein identification and quantification

Protein identification and quantification was performed using MaxQuant version 1.5.3.30 [73, 74]. The data were searched against the *Homo sapiens* UniProt database [75]. For shotgun SILAC experiments, the following parameters were used: peptide mass accuracy, 20 parts per million (ppm) for first search, 10 ppm for second search; trypsin enzyme specificity, fixed modifications, carbamidomethyl; variable modifications, methionine oxidation, deamidation (NQ), and N-acetylation (protein N-terminus); and all other parameters as preset. For SEC-PCP-SILAC experiments, the following parameters were used: peptide mass accuracy, 10 ppm; fragment mass accuracy, 0.05 Da; trypsin enzyme specificity; fixed modifications, carbamidomethyl; variable modifications, methionine oxidation, deamidation (NQ), and N-acetylation (protein N-terminus). For shotgun SILAC and SEC-PCP-SILAC experiments, both the requantify and match between runs options were enabled. For SEC-PCP-SILAC experiments, SILAC labels with a minimum ratio count of one were included. For immunoprecipitation experiments, label-free quantitation (LFQ) was enabled, with a minimum ratio count of two. For analysis of purified ribosome composition, LFQ was enabled with a minimum ratio count of one, due to the small size of ribosomal proteins. Only those peptides exceeding the individually calculated 99% confidence limit (as opposed to the average limit for the whole experiment) were considered as accurately identified. In all analyses, contaminants and reverse hits were removed using Perseus version 1.6.1.1 [76].

### Data analysis for shotgun proteomics of the IFN response

Differentially expressed proteins were identified using the one-sample moderated t-tests implemented in limma [77], followed by Benjamini-Hochberg correction. Enriched Gene Ontology terms were identified for proteins with significant differential expression between conditions at 5% FDR using the conditional hypergeometric test [78] implemented in the GOstats R package [79], with terms from each branch of the ontology analyzed separately. Enrichment for protein-protein interactions between differentially expressed proteins was assessed using the InBioMap database [22]. The likelihood of the observed number of interactions between differentially expressed proteins was evaluated by randomly rewiring the InBioMap interactome 100 times using a degree-preserving algorithm [23] in order to maintain network topology, with the number of iterations for the edge rewiring algorithm set to 6.9 × the number of edges in each network [80]. Network analyses were implemented in the R package ‘igraph’.

### Removal of high-magnitude errors in protein quantitation

Errors in quantitation of either the heavy or light channel can lead to the introduction of spurious outliers into PCP-SILAC chromatograms. To minimize the impact of these outliers on downstream analysis of the PCP chromatogram matrices, we applied a recently developed algorithm, “modern” (model-free outlier detection for robust networks), to remove outliers prior to network reconstruction(Skinnider et al., unpublished data). The motivating assumption that underlies modern is that a single observation should not globally rewire the interaction profile of a protein across the entire network. The interaction profile of each protein is quantified in modern as the vector of Pearson correlation coefficients between that protein and all other proteins in the network. Each point in the chromatogram is removed in turn, and the interaction profile is recalculated upon removal of each point. The correlation between the original interaction profile and the interaction profile with a single point removed is defined as the autocorrelation. Autocorrelation statistics are converted to z scores, and outliers are defined as observations associated with an autocorrelation z-score less than the normal distribution z-score corresponding to a two-tailed family-wise error rate of 0.05, given the total number of points observed. Application of modern to the six PCP chromatogram matrices led to the removal of 1,308 of 245,841 protein quantifications (0.53%).

### Protein-protein interaction network reconstruction

High-confidence protein-protein interaction networks were reconstructed from raw SEC-PCP-SILAC profiles using PrInCE [29], our open-source pipeline for co-fractionation data analysis, available at https://github.com/FosterLab/PrInCE. PrInCE first implements basic data cleaning and filtering functionality to restrict analysis to high-quality chromatograms. Briefly, single missing values are imputed as the mean of the two neighboring points, proteins with fewer than five observations are removed, and a sliding average with a width of five fractions is used to smooth the chromatogram. PrInCE then fits a mixture of one to five Gaussians to the smoothed chromatograms, performs model selection using the bias-corrected Akaike information criterion [81], and discards chromatograms that could not be fit by a mixture of Gaussians. A machine-learning procedure is then used to assign an interaction score to each pair of co-fractionation profiles using a naive Bayes classifier. Features used by the classifier include the Pearson correlation of the raw chromatograms and its corresponding P-value, the Euclidean distance between the raw chromatograms, the Pearson correlation of the smoothed chromatograms, the number of fractions separating the maximal values of each chromatogram, and the smallest Euclidean distance between any pair of fitted Gaussians. PrInCE concatenates features from each replicate but processes each isotope channel separately, providing a total of eighteen features to two naive Bayes classifiers. Importantly, unlike other published approaches, all features provided to the classifier are derived solely from the data, and do not incorporate prior biological knowledge, providing greater power for novel interaction discovery [34].

In addition to the features calculated from the co-fractionation data, the naive Bayes classifier requires sets of true positive (TP) and false positive (FP) interactions as training data. Previously, we have observed that many known protein-protein interactions catalogued in literature-curated databases are highly assay- or context-specific, and developed a “universal gold standard” subset of the CORUM database [82] tailored to predicting interactions from co-fractionation data [83]. This subset was used to train the classifier, with intra-complex interactions considered true positives and inter-complex interactions considered true negatives, using ten-fold cross-validation and taking the medium of all folds as the final interaction score. The naive Bayes classifier assigns an interaction probability to every pair, and returns a ranked list with putative interactions ordered by their interaction probability. The precision of the network was calculated at each point in the ranked list as the ratio of true positives to true positives and true negatives among interactions at that probability or higher, and we retained control and stimulated networks at 70% precision.

### Validation of the biological relevance of the IFN Interactome

To validate the biological relevance of the protein-protein interaction networks recovered by PrInCE, we assembled an aggregate network including unique interactions found in either the medium or heavy isotopologue channels at 70% precision, and compared the aggregate network to randomly rewired networks using a degree-preserving algorithm as described above. Human Gene Ontology annotations [84] were obtained from the UniProt-GOA database [85] and processed with the R package ‘ontologyIndex’ [86]. Annotations with the evidence codes “IPI” (inferred from physical interaction), “IEA” (inferred from electronic annotation), “NAS” (non-traceable author statement), or “ND” (no biological data), or the qualifier “NOT,” were removed, and annotations were propagated up the GO hierarchy. Very broad GO terms (annotated to more than 100 proteins) were removed, and the total proportion of interacting protein pairs sharing at least one GO term in each ontological category (biological process, cellular compartment, molecular function) was calculated for both rewired and observed networks. The distribution of interacting pairs sharing at least one GO term in each category among randomized networks was used to calculate an empirical P value for the observed enrichment. The same procedure was followed to evaluate the tendency of interacting protein pairs to be associated with the same disease, using disease genes obtained from the Mouse Genome Database [87] and mapped to their human orthologs using InParanoid [42]. The tendency of interacting protein pairs to contain domains known to physically interact in a high-resolution three-dimensional structure was similarly assessed using domain-domain interactions obtained from the 3did database [35], with Pfam domain annotations obtained from UniPro t[75]. Coexpression of protein pairs was calculated using the tissue proteome dataset described by [88]. Phylogenetic profiles were constructed using the InParanoid database([42], with the similarity in phylogenetic profile of a protein pair defined as the Pearson correlation between the binary presence/absence vectors of each protein across all species [89]. To quantify the proportion of novel interactions in the stimulated vs. unstimulated conditions, known interactions were compiled from sixteen databases, including BIND [90], BioGRID [91], DIP [92], HINT [93], HIPPIE [94], HPRD [95], IID [96], InBioMap [22], MatrixDB [97], Mentha [98], MINT [99], MPPI [100], NetPath [101], PINA [102], Reactome [103], and WikiPathways [104]. From pathway databases, only the subset of information cataloguing physical PPIs was retained. Gene and protein identifiers used to catalog interactions in these databases were mapped to UniProt accessions using identifier mapping files distributed by UniProt.

### Functional analysis of the IFN interactome

To quantify functional differences between the IFN-stimulated and unstimulated interactomes, we developed a permutation-based statistical test at the network level (Figure S4A). Briefly, for each term in the Gene Ontology, we identified all proteins annotated with that term, then calculated the total number of interactions in the stimulated and unstimulated networks between these proteins, and the difference between the two networks. We then randomly rewired both control and stimulated networks 1,000 times using a degree-preserving algorithm and calculated the difference in randomized networks. The null distribution of the randomized networks was used to calculate a z-score, which was subsequently converted to a probability and adjusted for multiple hypothesis testing using the method of Benjamini and Hochberg. GO terms with statistically significant differences between stimulated and unstimulated networks at 10% FDR were visualized as an enrichment map [47] using the ‘ggnetwork’ package [105] and the Fruchterman–Reingold layout algorithm [106], as implemented in ‘igraph’. Edges were drawn between GO terms on the basis of the complete set of proteins each was annotated to if the Jaccard index was greater than 0.33. For select terms, we additionally plotted the distribution of differences in number of interactions in randomized networks as a histogram. Autocorrelations were calculated as the Pearson correlation between pairs of stimulated and unstimulated chromatograms, normalized to z scores, and aggregated across all three replicates using Stouffer’s method. dN/dS ratios between human and mouse genes were obtained from Ensembl BioMart [107]. The total proportion of species in which an ortholog of each gene was present was quantified using the InParanoid database [42]. pLI scores were obtained from ExAC [43] and quantile normalized.

### Evolutionary analysis of IFN-stimulated genes

Core, conserved and species-specific ISGs were obtained from Shaw et al. (2017), on the basis of upregulation in all vertebrates studied, in human and at least one other species, and in human only, respectively. The statistical significance of the difference between distributions of Pearson correlations of raw PCP chromatograms in the unstimulated and stimulated conditions was assessed using a Brunner–Munzel test; correlations based on fewer than five pairwise observations were excluded from analysis. Co-immunoprecipitations from stimulated and unstimulated cells were compared using limma [77], and enrichment for different classes ISGs in immunoprecipitations of stimulated cells was assessed by gene set enrichment analysis [108], as implemented in the R package ‘fgsea’ [109].

### Analysis of siRPL28 knockdown data

Time-course transcriptome profiles of mouse ISGs in CD19^+^ B lymphocytes were obtained from [8]. These profiles were normalized by maximum expression across all timepoints, as in the original analysis; mapped to their human orthologs using InParanoid [42]; and matched to proteins that were differentially expressed after 4 h or 24 h of IFN stimulation in our shotgun proteomics data. Analysis of the siRPL28 SILAC proteomics dataset required proteins to be quantified in both the heavy and medium channels in at least two of three replicates. Protein ratios were log-transformed and compared between siRPL28-treated and untreated, IFN-stimulated cells using the moderated t-test implemented in limma [77]. A consensus set of ISGs was defined as those upregulated in human and at least four other species (five of ten species total) in the Shaw et al. (2017) dataset, although the observed enrichment was insensitive to the precise number of species used to define this set. The enrichment for ISGs among up- or downregulated proteins was assessed by gene set enrichment analysis [108], as implemented in the R package ‘fgsea’ [109]. The abundance profiles of selected ISGs were plotted after normalization by subtraction of mean abundance and division by the standard deviation.

## Supplementary Figures

**Figure S1.**
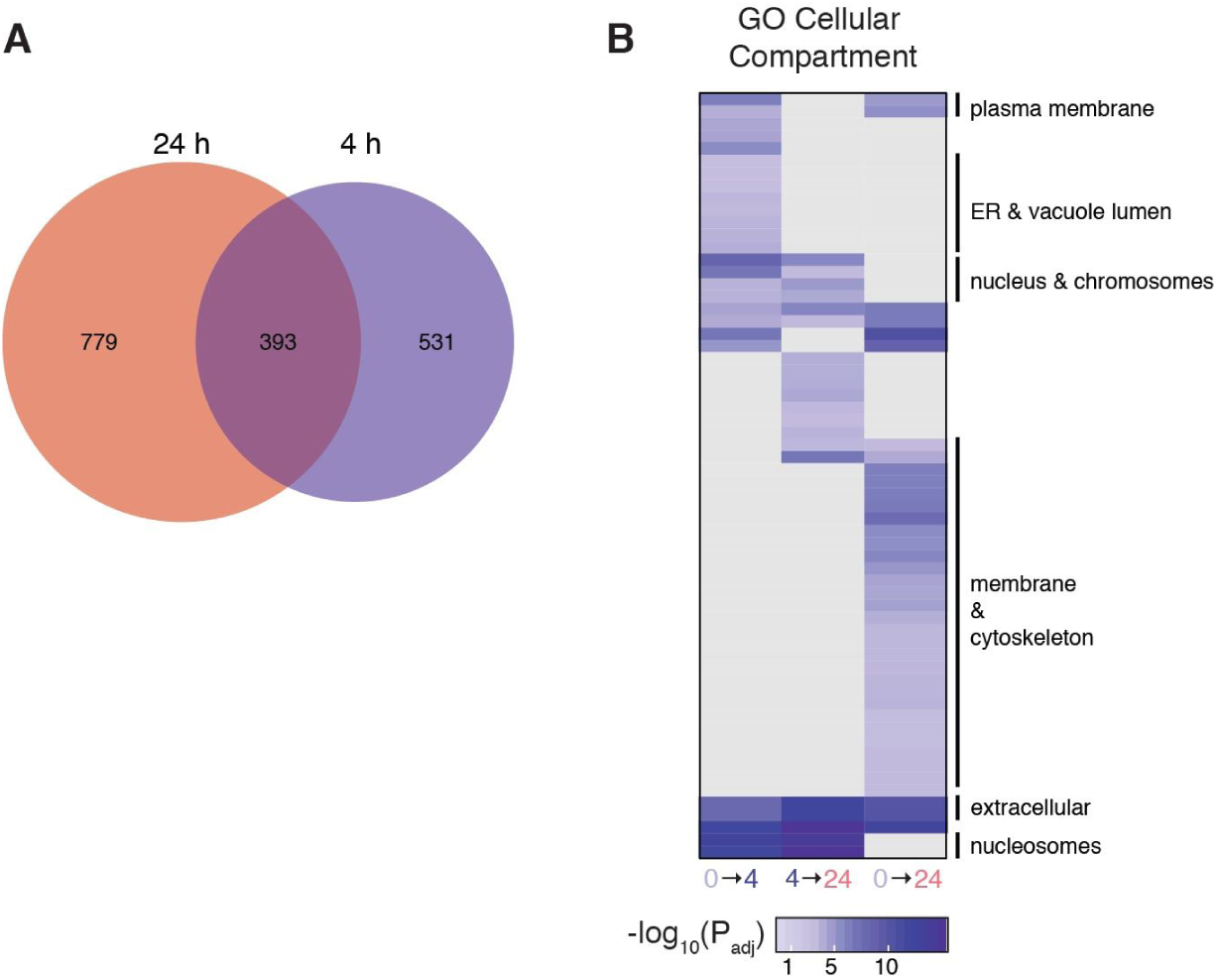
Proteomic characterization of the type I IFN response. (A) Venn diagram of overlap between differentially expressed proteins after 4 h and 24 h of IFNβ stimulation, relative to unstimulated cells. (B) Gene Ontology (GO) terms for cellular compartments significantly enriched among differentially expressed proteins after 4 h or 24 h of IFNβ stimulation.

**Figure S2.**
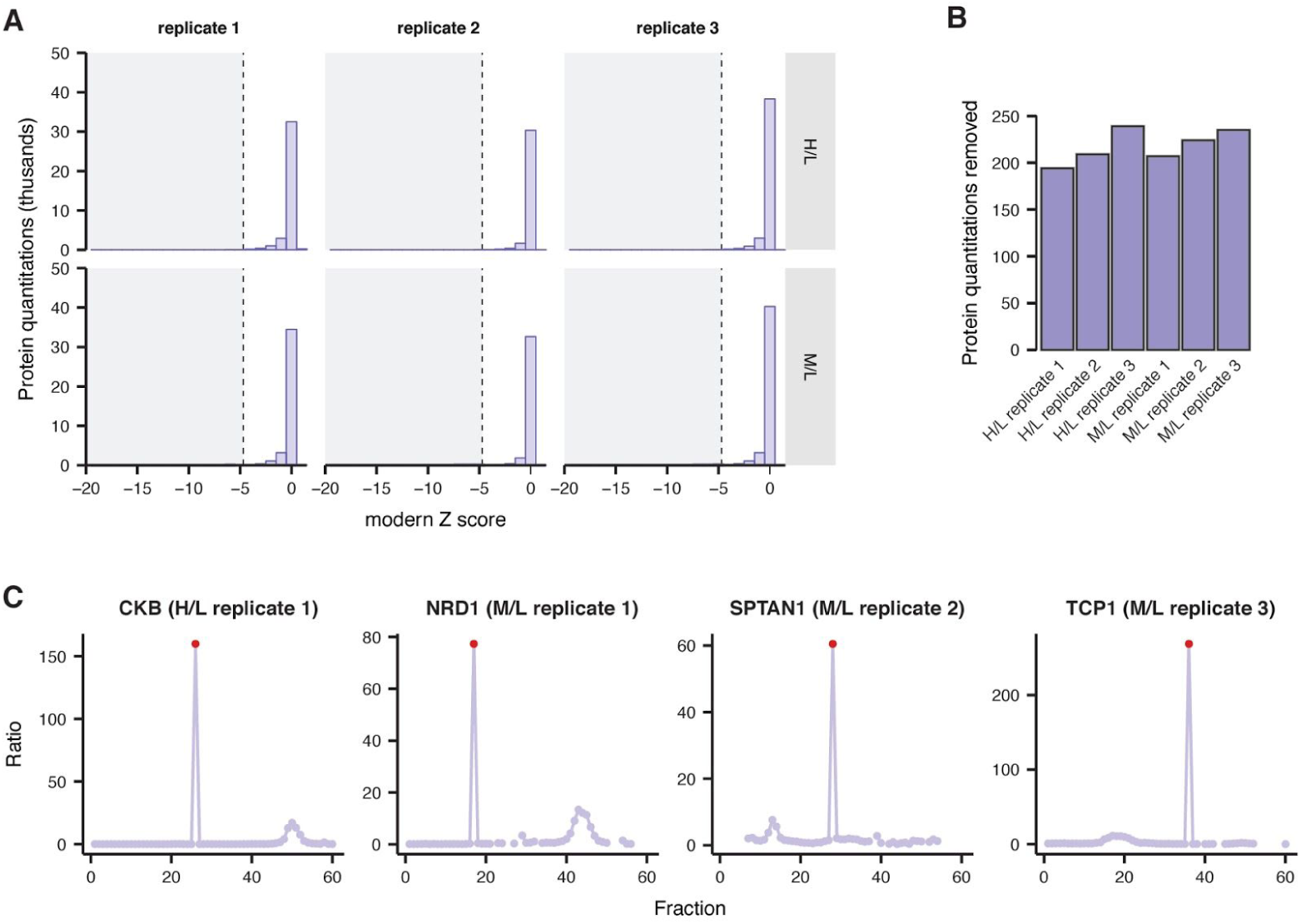
Removal of high-magnitude errors in protein quantitation prior to protein-protein interaction network inference by modern. (A) Distribution of z scores for 245,841 total protein quantitations across three biological replicates, as determined by modern (Methods). (B) Total number of protein quantitations censored in each biological replicate, based on an autocorrelation z-score less than the normal distribution z-score corresponding to a two-tailed family-wise error rate of 0.05, given the total number of points observed. (C) Examples of high-magnitude errors in protein quantitation correctly detected by modern.

**Figure S3.**
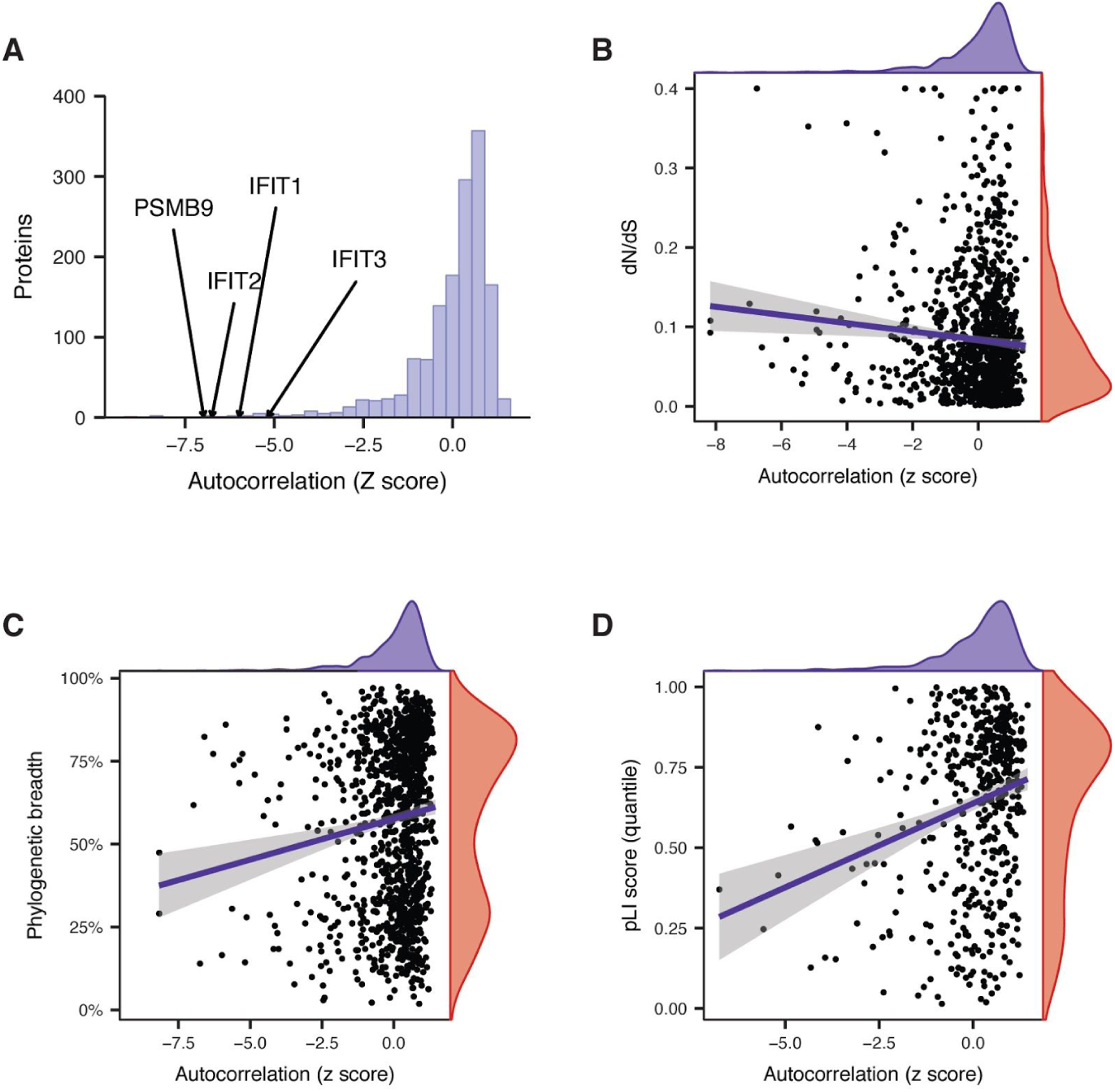
Evolutionary plasticity of the interactome response to IFN stimulation. (A) Distribution of autocorrelation z scores between IFN-stimulated and unstimulated protein correlation profiles, derived from Stouffer integration across all three replicates. Selected highly rewired proteins are highlighted. (B) Relationship between autocorrelation z score and ratio of nonsynonymous to synonymous substitutions (dN/dS; (Spearman ρ = –0.11, p = 1.9 × 10^−4^). (C) Relationship between autocorrelation z score and proportion of genomes in the InParanoid database [42] in which an ortholog of the protein was present ρ = 0.14, p = 5.3 × 10^−7^). (D) Relationship between autocorrelation z score and pLI score quantile (Spearman ρ = 0.18, p = 5.9 × 10^−5^).

**Figure S4.**
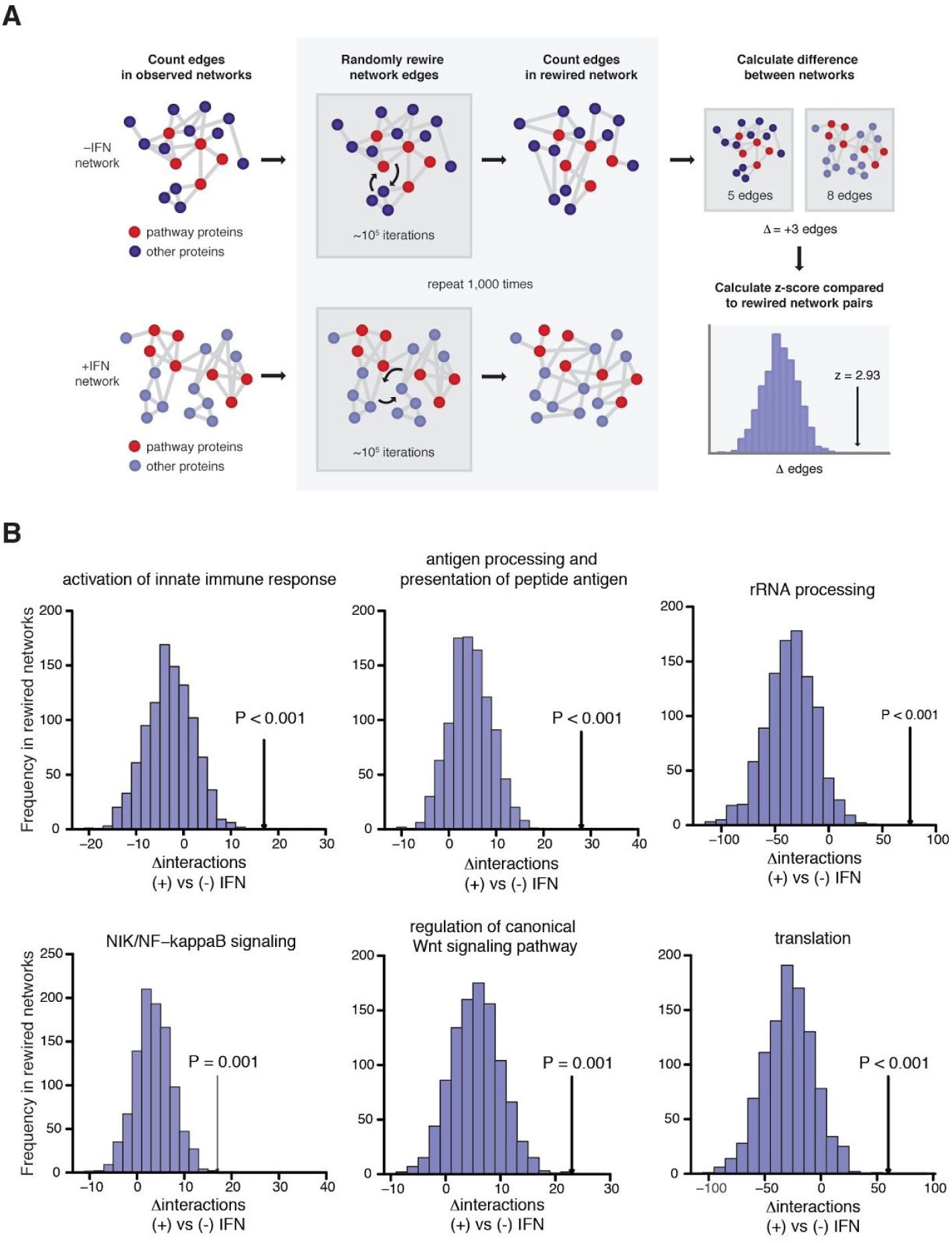
A statistical framework for differential network analysis at the functional level. (A) Schematic overview of the workflow for functional differential network analysis (Methods). For a given Gene Ontology (GO), the total number of interactions between proteins annotated with that term was calculated in stimulated and unstimulated networks. The difference in the number of interactions involving that term between conditions was then compared to the difference between 1,000 randomly rewired versions of the same networks. (B) Examples of GO terms significantly enriched in the IFN-stimulated network. Histograms show the difference in the number of interactions between 1,000 randomly rewired versions of the IFN-stimulated and unstimulated networks; arrows show the observed difference in the number of interactions.

**Figure S5.**
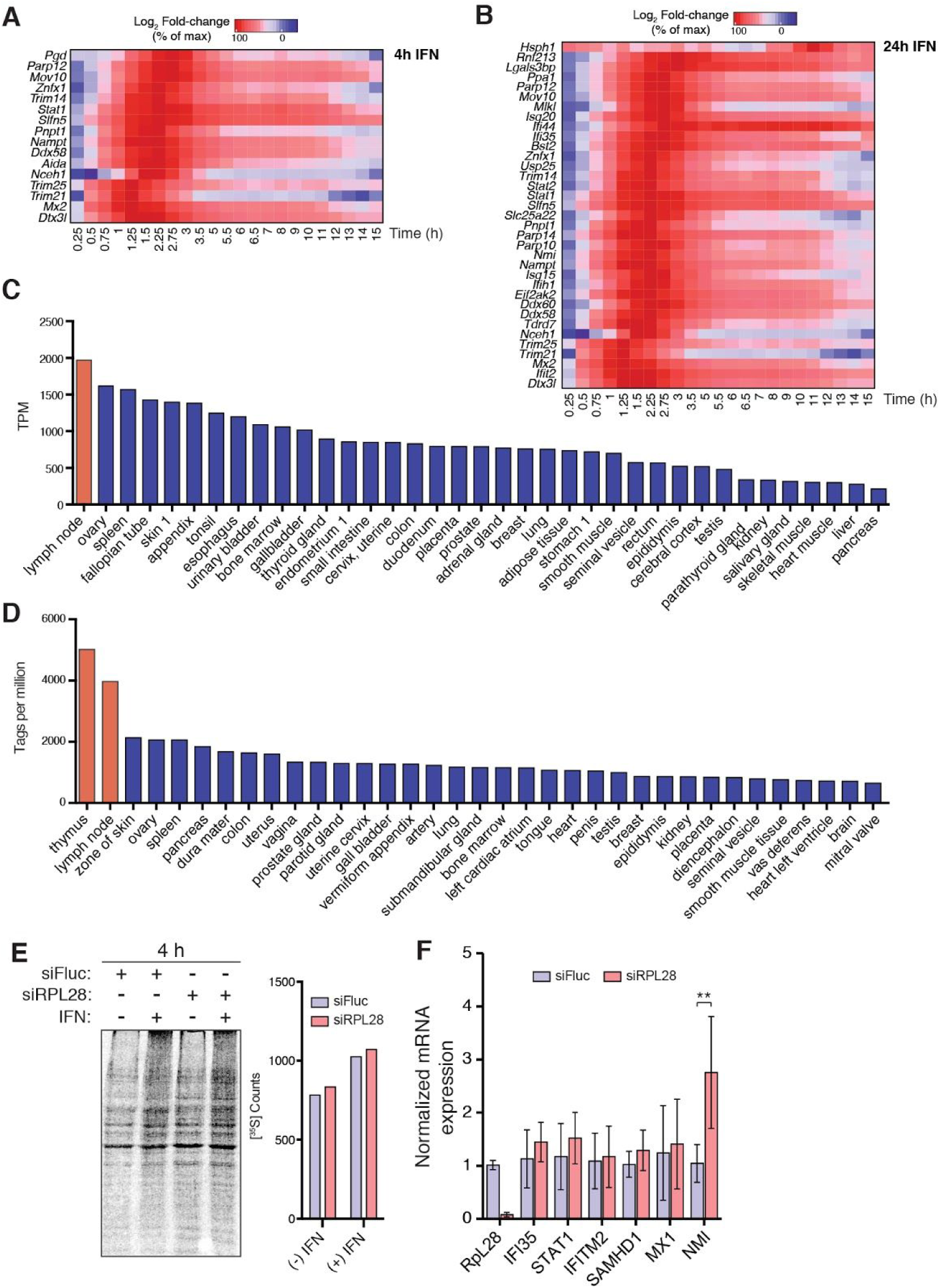
A specialized translational program mediates the effects of IFN stimulation on protein synthesis. (A–B) mRNA expression profiles of proteins that were significantly upregulated at the proteomic level after 4 h (A) or 24 h (B) of IFN stimulation in fine-grained time-course data [8]. (C) RPL28 mRNA expression in human tissues from the Human Protein Atlas [61]. (D) RPL28 mRNA expression in top 35 human tissues from the FANTOM5 database. (E) Metabolic labeling of HeLa cells by [^35^S]-Met/Cys incorporation after siRNA knockdown of RPL28 for 48 h followed by IFN stimulation. Left: representative gel of [^35^S]-Met/Cys labeled proteins at 4 h post-IFN treatment. Right: average quantitation of [^35^S] counts in precipitated protein from two independent experiments at 8 h post-IFN treatment. (F) qRT-PCR of various ISGs from cells treated with siRPL28 or control siRNA for 48 h followed by IFN treatment for 8 h. Expression levels were normalized to *TUBB* then to control. ** *P* < 0.005.

## SUPPLEMENTARY TABLES

**Table S1:** Shotgun proteomic SILAC ratios of IFN-stimulated cells at 4 h or 24 h IFN treatment.

**Table S2:** GO term enrichment on significantly changed SILAC ratios of IFN-stimulated cells at 4 h or 24 h IFN treatment.

**Table S3:** SEC-PCP-SILAC interactomes of unstimulated and IFN-stimulated HeLa cells at 70% precision.

**Table S4:** Differential interactions between (-) and (+) IFN.

**Table S5:** Label-free quantitation intensities of ribosomal proteins.

**Table S6:** Shotgun proteomic SILAC ratios of siRPL28-treated cells plus IFN for 8 h.

